# Investigating the ability of deep learning-based structure prediction to extrapolate and/or enrich the set of antibody CDR canonical forms

**DOI:** 10.1101/2023.12.08.570786

**Authors:** Alexander Greenshields-Watson, Brennan Abanades, Charlotte M Deane

## Abstract

Deep learning models have been shown to accurately predict protein structure from sequence, allowing researchers to explore protein space from the structural viewpoint. In this paper we explore whether “novel” features, such as distinct loop conformations can arise from these predictions despite not being present in the training data.

Here we have used ABodyBuilder2, a deep learning antibody structure predictor, to predict the structures of ∼1.5M paired antibody sequences. We examined the predicted structures of the canonical CDR loops and found that most of these predictions fall into the already described CDR canonical form structural space. We also found a small number of “new” canonical clusters composed of heterogeneous sequences united by a common sequence motif and loop conformation. Analysis of these novel clusters showed their origins to be either shapes seen in the training data at very low frequency or shapes seen at high frequency but at a shorter sequence length.

To evaluate explicitly the ability of ABodyBuilder2 to extrapolate, we retrained several models whilst withholding all antibody structures of a specific CDR loop length or canonical form. These “starved” models showed evidence of generalisation across CDRs of different lengths, but they did not extrapolate to loop conformations which were highly distinct from those present in the training data. However, the models were able to accurately predict a canonical form even if only a very small number of examples of that shape were in the training data.

Our results suggest that deep learning protein structure prediction methods are unable to make completely out-of-domain predictions for CDR loops. However, in our analysis we also found that even minimal amounts of data of a structural shape allow the method to recover its original predictive abilities. We have made the ∼1.5 M predicted structures used in this study available to download at https://doi.org/10.5281/zenodo.10280181.

## Introduction

Deep learning has revolutionised the field of structural biology with tools such as AlphaFold2 (AF2) (Jumper et al., 2021), RosettaFold (Baek et al., 2021) and ESM-Fold (Lin et al., 2023) that can accurately predict protein tertiary structure from primary sequence. These tools are all trained on the known protein structure landscape derived from the PDB (Berman et al., 2000) and have been shown to generalise well to proteins that were not seen during training. Several studies have used these models to enrich the existing protein structure landscape by making extensive predictions from the larger available sequence space. Analysis of these predictions revealed many examples of structures that are very different from the closest available match in experimentally defined data (Bordin et al., 2023; Lin et al., 2023).

By analysing over 365,000 high confidence structures predicted by AF2, Bordin et al. were able to define 25 novel superfamilies which did not cluster into any existing CATH classifications using their CATH-Assign protocol (Bordin et al., 2023). A second example of new knowledge arising from structural predictions was provided by ESMFold (Lin et al., 2023). Here, Lin et al. predicted the structures of over 600M metagenomic sequences isolated from diverse environmental and clinical samples. The use of these metagenomic sequences increased the probability of finding examples that were highly distant from the sequence and structural data used to train ESM2 and ESMFold respectively (Lin et al., 2023). Within a sample of 1M modelled structures defined as high confidence (predicted local distance difference test score, pLDDT *>* 0.7 and predicted template modelling score, pTM *>* 0.7), the authors found over 125,000 predictions with no close match in the PDB (defined as pTM *>* 0.5 carried out using Foldseek (van Kempen et al., 2023)) and in close alignment to the corresponding predictions from AF2. While both studies demonstrate that structure prediction tools can confidently generate novel structures, X-ray crystallography data was not obtained to conclusively validate the predictions. It is also not clear if the novel structures generated are composites of large substructural fragments present in the training data.

To attempt to explicitly address whether models can generalise to unseen regions of structural space, Ahdritz et al. carried out ‘out-of-domain’ experiments using OpenFold (Ahdritz et al., 2022). In particular, examining if OpenFold can generalise from limited data to accurately predict alpha helices or beta sheets despite their omission from training datasets. However, they were not able to completely remove all signal of these secondary structures from their training data, and hence the models were likely still learning from a much-reduced set of examples, rather than extrapolating to a completely unknown structure based on their induction of biophysical rules.

These analyses raise the question of whether current deep learning-based models are truly capable of predicting conformations which are never present in training data. While extrapolation by deep neural networks is theoretically plausible (Balestriero et al., 2021; Fannjiang and Listgarten, 2023) searching for evidence of this is difficult and requires extensive classification of training data and the resulting predictions.

One limitation of deep learning based protein structure predictors is their poor performance on stretches of sequence that are intrinsically disordered (Ruff and Pappu, 2021; Tunyasuvunakool et al., 2021) or explore diverse conformational space (Chakravarty and Porter, 2022). The loops of adaptive immune receptors, antibodies, and T cell receptors, fall into the latter category. These loops form the majority of the binding site (paratope) of these proteins and are termed complementarity determining regions (CDRs) (Schatz and Ji, 2011).

The protein sequences that make up the CDR loops arise from two genetic mechanisms, termed V(D)J recombination (Alt et al., 1984; Brack et al., 1978) and somatic hypermutation (Griffiths et al., 1984). In antibodies, the process of V(D)J recombination randomly pairs V-, D-and J-genes (VJ genes for light chains and VDJ genes for heavy chains) and introduces junctional diversity through insertion and deletion of nucleotides. Further diversity is introduced to the V-gene region of the antibody by somatic hypermutation, where point mutations that modify the amino acid sequence and improve binding affinity are positively selected and progressively dominate the immune response to a pathogen. These mechanisms create high levels of sequence diversity and are evolutionarily advantageous as they combine to provide a nearly limitless potential of binding solutions which allow antibodies to neutralise the correspondingly limitless diversity of pathogens to which humans can be exposed (Laserson et al., 2014). Of the six CDR loops on an antibody, diversity is highest within the CDRH3, the residues of which often disproportionally govern paratope-epitope interactions (Regep et al., 2017). Structure predictors have been found to perform poorly on this region of antibodies, for example with average RMSD values between predictions and ground truth that exceed 2.5 Å for state-of-the-art models such as AlphaFold-Multimer (AFM) (Evans et al., 2022) and ABody-Builder2 (ABB2) (Abanades et al., 2023).

The predictive performance on the remaining five loops, CDRL1-3 and CDRH1-2, is far better (average RMSD *<* 1Å), despite these being subject to the genetic process of somatic hypermutation and being influenced by neighbouring hypervariable loops (Guloglu and Deane, 2023). The ability to accurately model these can be explained by canonical forms, the term given to sets of CDR loops of the same length that adopt similar backbone conformations and share a sequence motif (Chothia et al., 1989). These canonical forms were first observed in crystallographic datasets of available antibody structures before 1986 (Chothia and Lesk, 1987). With the deposition of more structural data both the number of canonical conformations and the sequences that could be assigned to each were continuously expanded and redefined (Adolf-Bryfogle et al., 2015; Kelow et al., 2022; North et al., 2011; Nowak et al., 2016; Wong et al., 2019a, 2019b). This information linked diverse sets of sequences to distinct loop conformations and thus was increasingly useful to antibody researchers, by providing a form of sequence-to-structure prediction that could be automated by template search and homology modelling tools (Sivasubramanian et al., 2009; Wong et al., 2019a).

The latest CDR structure and sequence pairings harvested from antibody structural data are defined in PyIgClassify2 (Kelow et al., 2022). The definitions were released as the ‘penultimate classification of canonical forms’ in reference to the breakthroughs in structure prediction research that may soon render the predictive power of this relationship obsolete. While structure prediction methods are still being evaluated, especially in the domain of adaptive immune receptors, PyIgClassify2 can serve as a map of the known conformational space explored by antibody CDRs.

Using the rigorous definitions from PyIgClassify2 as a reference point in structural space, we set out to test whether the predicted structures from all available paired antibody sequences in observed antibody space, OAS, (Kovaltsuk et al., 2018; Olsen et al., 2022) reveal novel canonical clusters or highlight conformations not explored by existing experimental data. We then assess these new areas to determine whether they represent evidence of extrapolation or had direct origins in the training data.

ABodyBuilder2 (ABB2) is a structure prediction tool specific to antibodies (Abanades et al., 2023). It uses an ensemble of four deep learning models trained on the structures of over 3500 antibodies as well as a fast minimisation in the AMBER14 forcefield (Eastman et al., 2017; Maier et al., 2015) to make predictions with comparable accuracy to AFM in a fraction of the time. We used ABB2 to predict the structures of ∼1.5M paired antibody sequences. We mapped the conformational space of the CDRL1-3 and CDRH1-2 loops and used existing classifications of the canonical forms in experimental data as reference points. By comparing the loop conformations of canonical clusters to clusters found in predictions derived from the ∼1.5M heterogeneous sequences we were able to redefine and identify new canonical clusters.

These new clusters (potential canonical forms) were defined by unique sequence motifs and shared loop conformations and typically arose from enrichment of a small number of examples in the experimental data. We also observed apparently novel clusters (canonical forms) that derived from similar shapes (and sequences) of a different loop length, a phenomenon that has been previously described within the structural dataset (Nowak et al., 2016), termed length independence.

Using our mapping of structural space and the definitions of canonical forms we designed out-of-domain retraining experiments which explicitly tested the capability of ABB2 to both generalise and extrapolate. We found that with zero examples of a given CDR shape ABB2 was consistently unable to predict it. However, with the introduction of very small numbers of a shape, the predictive ability was restored. Overall, these analyses exemplify the power of augmenting experimental data with predictions and provide simple tests for extrapolation and effective data recapitulation that may help inform the next generation of structure predictors.

## Method

### Selection of paired antibodies sequences for ABodyBuilder2

Paired antibody sequences were retrieved from observed antibody space (OAS) (Kovaltsuk et al., 2018; Olsen et al., 2022) (1-March-2023, https://opig.stats.ox.ac.uk/webapps/oas/). Non-redundant sequence pairs were filtered by minimum lengths defined in the ABB2 workflow such as minimum residue length of 70, starting IMGT residue number less than 8, and end residue number greater than 120, (see sequence checks https://github.com/oxpig/ImmuneBuilder). Sequences containing any gaps or ambiguities were removed to leave 1,492,044 pairs for processing. ABB2 was run on all sequences and 1,492,031 structures were successfully predicted.

### Annotation of CDR loops in paired antibody sequences

To group the relevant modelled structures for conformational analyses, the CDR loops of all input sequences were annotated with information on their sequence composition and length. The CDRs were defined according to the IMGT numbering scheme (CDR1: 27-38, CDR2: 56-65, CDR3: 105-117) (Lefranc and Lefranc, 2023). This numbering system was chosen as the anchor residues and CDR locations are consistent for both heavy and light chains, as well as being the standard reference point for V(D)J gene annotation. Each sequence was IMGT numbered using ANARCI (Dunbar and Deane, 2016) and the CDR sequences checked against corresponding information from IgBLAST annotations (Ye et al., 2013). For a given heavy or light chain, if there was any discrepancy between IgBLAST and ANARCI CDR definitions then this chain was not taken forward for conformational analysis (resulting in the exclusion of 6414 light chains and 9822 heavy chains from the structures predicted above).

### Retrieval and selection of experimental structures from SAbDab

The structural antibody database (SAbDab) (Dunbar et al., 2014; Schneider et al., 2022) is a curated database which contains all antibody, single chain variable fragment (SCFV) and nanobody structures available in the PDB (Berman et al., 2000). IMGT numbered structures used in ABB2 test, train and validation datasets were downloaded from SAbDab. These structures were derived by X-ray crystallography or cryogenic electron microscopy (cryo-EM) and with a resolution better than 3.5 Å (full list given in SI of Abanades et al. 2023).

### Annotation of SAbDab structures with PyIgClassify2 information

The information on CDR loop canonical forms was obtained from the pyig cdr data.txt file downloaded from the PyIgClassify2 website (http://dunbrack2.fccc.edu/PyIgClassify2/, 21-Feb-2023). This provides complete information for all CDR loops on each structure within a given PDB file, including information on sequence identical but structurally distinct members of the asymmetric unit which are distinguished by their PDB chain identifier. For each loop the relevant information includes the length, sequence, canonical form assignment (both with and without an electron density confidence cut off), PDB identifier of the parent chain and information on whether the CDR is structurally complete or is missing any backbone coordinates. This data was used to filter the relevant structures and their CDR loops for each analysis, and to annotate the experimental data points by canonical cluster membership.

### Alignment of IMGT and AHo numbering systems

PyIgClassify2 CDR lengths are defined according to the AHo numbering scheme (Honegger and Plückthun, 2001) which symmetrically places insertions and deletions around positions defined as key residues in each CDR. This deviates from the IMGT numbering scheme (Lefranc and Lefranc, 2023) which places insertions centrally within each CDR at fixed positions. The different approaches to defining the CDRs mean that IMGT defined lengths for CDRL1-2 and CDRH1-2 are shorter than those defined in the AHo numbering scheme used in PyIgClassify2. Therefore, for each CDR and length combination analysed in this study, we have listed the corresponding AHo CDR lengths in Table 1, along with the PyIgClassify2 defined canonical forms and sequence motifs described in (Kelow et al. 2023).

**Table 1:**
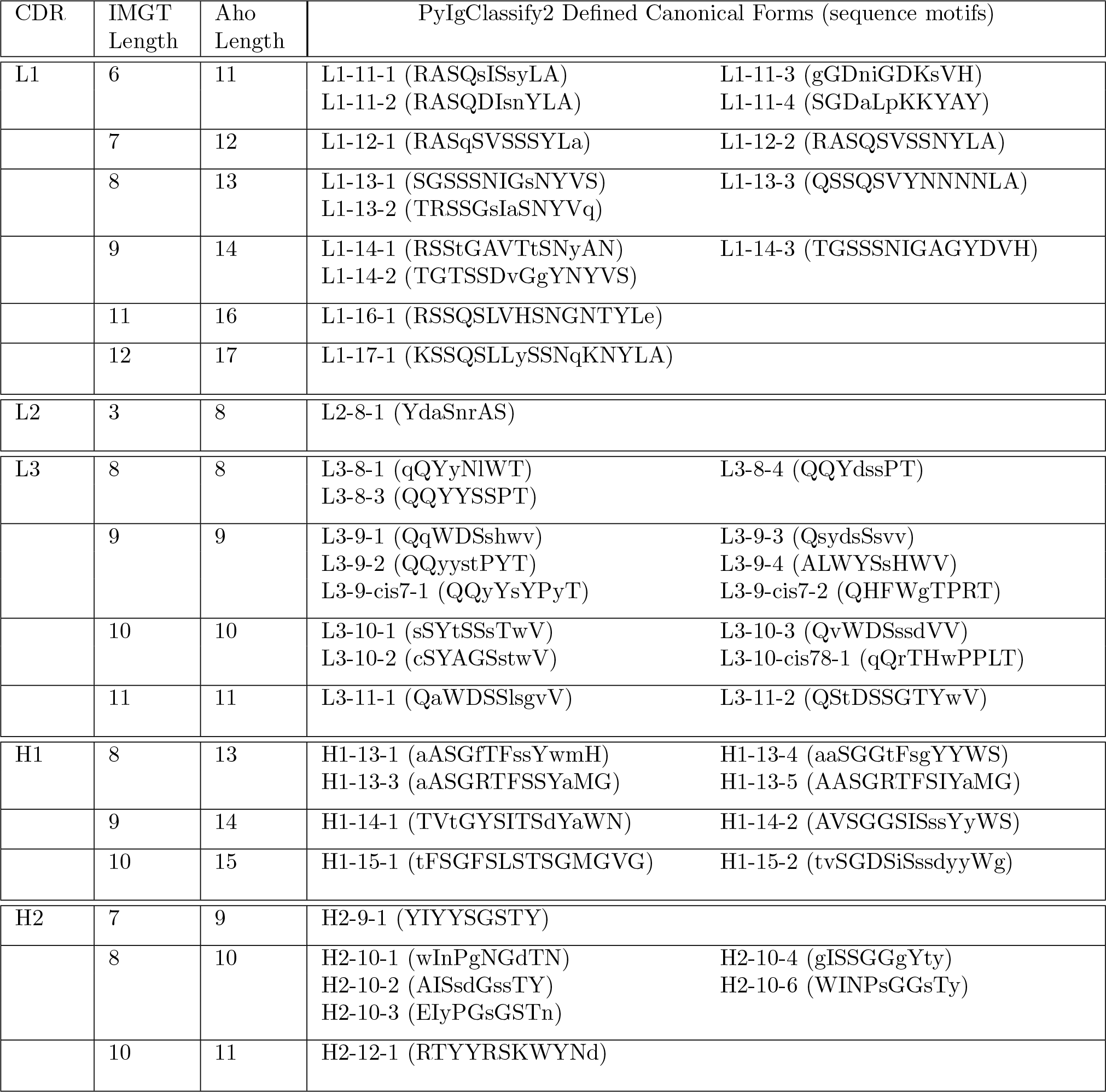
Alignment of IGMT CDR numbering, AHo CDR numbering and PyIgClassify2 Canonical Forms. For each CDR, the IMGT lengths used in these analyses are presented alongside the corresponding AHo lengths and PyIgClassify2 defined canonical forms for that CDR. PyIgClassify2 canonical forms are named according to the CDR (e.g. L1) followed by the Aho length (e.g. -6), and the form number (e.g. -1) to form a unique identifier for each form (e.g. L1-11-6). For each canonical form the consensus sequence motif, as defined in (Kelow et al. 2022) is given in brackets. Uppercase letters of the consensus sequence indicate highly conserved amino acids at a given position, while lowercase letters indicate a less conserved amino acid that was still observed in the cluster of loops found in the PyIgClassify2 analyses. Information is only provided for the loops analysed in this study, i.e., those which corresponded to the most dominant non-redundant sequences in OAS, (see Fig. 1B).

### Pre-processing of SAbDab datasets

SAbDab files included SCFV structures where the heavy and light chain are part of a single continuous sequence, as well as datasets with multiple sequence-identical copies in the asymmetric unit. To ensure all CDR loops could be correctly identified and consistently aligned, the relevant chains in each dataset were isolated, IMGT numbered and then saved as individual files linked to the corresponding PyIgClassify2 meta data. For SCFV structures the continuous sequence was broken and each fragment treated as an individual chain. If a chain within a dataset could not be numbered with ANARCI, or a specific CDR loop was missing residues (as indicated by the PyIgClassify2 ‘cdr ordered’ flag), they were not included in structural analyses. This processing resulted in 11821 heavy and light chains from 3355 PDB files which were used for further analyses. As ABB2 predicted structures were all correctly numbered and contained only a single copy in the asymmetric unit, they did not require any preprocessing for downstream analyses.

### Structural analysis of CDR loops of the same length

To perform structural analysis, CDR loops of predicted structures were grouped according to their CDR type (i.e., CDRL1 or CDRH2), amino acid length and sequence composition (non-redundant sequences only). These were analysed alongside all relevant loop structures from SAbDab, for these experimental data points redundant sequences were included as they may contain alternate conformations of the same sequence.

To provide a consistent frame of reference for each CDR length, a loop template was chosen from the highest resolution PDB structure available, this structure also had to be classified as representative of a PyIgClassify2 defined canonical form (‘is representative’ flag) and thus was not likely to be an outlier or exhibit any structural features that set it apart. All CDR loops in the analysis were aligned to this template by superimposition of the alpha carbon atoms of the 10 framework residues either side of the loop (CDR1: 22-26 & 39-43, CDR2: 51-55 & 66-70, CDR3: 100-104 & 118-122). If superimposition resulted in RMSD values greater than 1.5 Å then these were not taken forward for loop comparisons. For predicted structures, the framework regions were highly consistent and less than 5 loops per analysis were eliminated. For experimental data points the number eliminated due to framework misalignments ranged from 0 to a maximum of 31 for CDRL1-Len-6 (out of 2499 chains), with a median number of 3 data points eliminated across all analyses. The carbon and nitrogen backbone atom coordinates of the aligned loops were extracted and saved (CDR1: 27-38, CDR2: 56-65, CDR3: 105-117). Atom counts were checked and then all pairwise RMSD values calculated. This resulted in an N-by-N pairwise distance matrix of RMSD values including both predicted and experimental datapoints for each CDR loop type at every length. To limit the size of pairwise matrices, loops of predicted structures were analysed in batches of 42,000. Where a CDR and length had more than this number of non-redundant sequences (see Table 2) subsequent batches of a maximum size of 42,000 were run until all relevant loop structures were analysed. All batches were run through the clustering and visualisation pipeline (see sections below) and then inspected to ensure results were consistent across all analyses. For multi-batch CDRs, graphs of the first batch are shown in main figures and graphs of subsequent batches are provided in the supplementary information (Supp Fig. 7).

**Table 2:** Results of RMSD and DBS cluster analysis in high frequency CDR lengths. For each CDR and IMGT length explored in this study, details of the number of non-redundant sequences (and hence loops structurally analysed) and the corresponding results of structural analyses are given. The new information arising from analysis of the predicted structures could be defined as either the identification of a new canonical cluster, or the sub-division of an existing cluster of loops that had previously been defined as belonging to a single canonical form. Further details are given on whether these clusters arose from length independent conformations and/or existing experimental data points classified as unassigned by PyIgClassify2 (Kelow et al. 2022).

| CDR | IMGT Length | Number of Unique Sequences in OAS | New information arising from predictions? | Origin of novel canonical cluster? |
| --- | --- | --- | --- | --- |
| L1 | 6 | 21,707 | No | N/A |
|  | 7 | 10,212 | New canonical cluster | Extra density matching existing unassigned exp data |
|  | 8 | 7,596 | No | N/A |
|  | 9 | 13,846 | New canonical cluster | Extra density matching existing unassigned exp data |
|  | 11 | 6,992 | Sub-division of existing cluster | Uneven distribution of density within existing form |
|  | 12 | 10,565 | No | N/A |
| L2 | 3 | 2,280 | No | N/A |
| L3 | 8 | 15,805 | No | N/A |
|  | 9 | 76,087 | No | N/A |
|  | 10 | 56,599 | New canonical cluster | Density derived from length-independent conformation and existing data |
|  | 11 | 51,277 | New canonical cluster | Density derived from length-independent conformation |
| H1 | 8 | 61,617 | Sub-division of existing cluster | Uneven distribution of density within existing form |
|  | 9 | 5,655 | No | N/A |
|  | 10 | 25,000 | Sub-division of existing cluster | Extra density matching existing unassigned exp data |
| H2 | 7 | 27,769 | Sub-division of existing cluster | Uneven distribution of density within existing form |
|  | 8 | 99,003 | No | N/A |
|  | 10 | 13,114 | No | N/A |

### Structural analysis of CDR loops of different length

For later analyses aimed at discovering length independent conformations, we also calculated the distance between loops of different lengths. The normalised dynamic time warping (DTW) scores were used to quantify the relative distance in groups of loops that included both length matched and mismatched pairs. Loops were aligned as described above to a high-resolution template, then raw DTW scores calculated between the coordinates using the ‘dtaidistance’ library in python (Meert et al., 2020). Normalisation was applied by squaring the score, dividing by the number of atoms being compared, and taking the square root (this resulted in a DTW score that is equivalent to RMSD when lengths are matched). The squared score was divided by the maximum number of atoms (i.e., for length 9 versus 8, the score was divided by 27 to account for 9 residues, each comprised of two carbon and one nitrogen backbone atoms).

### Density based clustering

The pairwise distance matrices of RMSD or DTW values contain information that represents the structural relationships between all loop conformations in an analysis. To identify clusters of loops with similar conformations within this high-dimensional data, we employed density-based clustering, using the density-based spatial clustering of applications with noise (DBSCAN) function from the scikit-learn library (SK learn) (Pedregosa et al., n.d.). The DBSCAN function takes the input distance matrix and two parameters: the minimum number of data points required to form a cluster (min points) and the minimum distance between any two points in the same cluster (epsilon). The number and size of clusters identified in each analysis is highly sensitive to these values and must be optimised based on the input data. Therefore, we systematically calculated these values in a consistent manner for all analyses, allowing us to find the maximum number of clusters within high-dimensional space and assess whether any cluster related to a novel canonical form. Min point values were calculated by taking the square root of either the number of loops being compared, or the same number divided by 2. For epsilon values we performed a K-nearest neighbours (KNN) analysis on the distance matrix for values of K ranging from 2 to 5. For each value of K, the elbow point was taken from a scree plot of all KNN distances, and this elbow point value used as epsilon in subsequent DBSCAN analyses. This resulted in four separate DBSCAN analyses (one for each value of K from 2 to 5). To select the most appropriate analysis for cluster inspection and visualisation, we matched the K value used to determine epsilon with the number of clusters identified by DBSCAN. If there was no match, then the next closest match was taken for the highest value of K. While dominant clusters were often evident and easily found using multiple values of epsilon, the impact of optimising these parameters was most apparent when identifying smaller or overlapping clusters. We call the clusters generated in this way DBS clusters (short for DBSCAN-based selection).

### Inspection of density-based cluster structures and sequences

We manually inspected the loops comprising each DBS cluster by visualisation of both structures and sequences. This allowed us to assess the structural difference between each cluster and relate the sequence logos back to the defined sequences of PyIgClassify2 canonical clusters. The aligned 3D loops that were assigned to each DBS cluster were visualised using PyMol (Delano, 2002). We selected random samples of up to 20 loops from the predicted structures, all of which belonged to a specific DBS cluster. Samples were coloured according to cluster membership and viewed in the same frame. These loops were presented in multiple orientations to highlight backbone differences that led to distinct cluster assignment. For sequence logo plots, sequences from the predicted structures which had been assigned to a DBS cluster were plotted in R using the ggseqlogo package (Wagih, 2017). Logo plots are shown in the bitwise format (opposed to the proportion format) to maximise identification of the dominant amino acid enrichments and motifs specific to each cluster.

### Multidimensional scaling visualisation

To simplify the complex high-dimensional pairwise distance matrix and allow for easy visualisation, we applied multidimensional scaling (MDS) to create a 2D representation. We used the parallelised MDS function from the ‘lmds’ R package, before processing and plotting the output data using tidyverse packages (Wickham et al., 2019). These plots were then annotated according to the DBS cluster membership of each data point, or the canonical cluster assignment of only the experimentally derived data points.

### ABodyBuilder2 out-of-domain experiments

Out-of-domain experiments involved the removal of all data points related to a specific CDR length, or a specific canonical cluster, from both the training and test datasets of ABB2. The model was then trained on this modified dataset from scratch. The criteria for removing data from ABB2 training samples involved dividing the MDS map into quadrants and selecting the quadrant with the most distinct canonical cluster, i.e. clearly separated from other data points. All experimental data points in this quadrant were excluded. To ensure the removal of all relevant data points from the training data, any samples defined as the excluded canonical form by PyIgClassify2 without an electron density cutoff (using the ‘cluster nocutoff’ flag) were also eliminated.

### Data inclusion experiments

For out-of-domain experiments where small numbers of the excluded canonical form were reintroduced into the training data, we first added the highest-resolution datasets identified as representative of the missing canonical form (determined by the ‘is representative’ flag in PyIgClassify2). Subsequently, we progressively reintroduced the next highest-resolution datasets not designated as representative but still belonging to the high-confidence canonical cluster.

### Retraining of ABodyBuilder2

The original ABodyBuilder2 model consists of an ensemble of four models each trained independently. To make a prediction the outputs from each model are averaged and the prediction closest to average is selected as the final output. Of the four models, one utilised a 128-dimensional embedding and three utilised a 256dimensional embedding. Models were trained until no further improvement was seen in the validation loss after 100 epochs.

To facilitate the training of multiple models, for the initial experiments in this study we retrained a single model with a 128-dimensional embedding (not an ensemble). Each model was trained on either a Nvidia GeForce GTX-1080 Ti GPU or Quadro RTX 6000/8000 GPUs for 150-340 epochs for each training stage (see training methods Abanades et. al 2023), continuing until no further improvement in validation loss was observed after 50 epochs (half the number used to train original ABB2 models). For specific retraining experiments (those used to confirm data inclusion thresholds important for prediction), an ensemble of models was created, each consisting of one model with a 128-dimensional embedding and three models with a 256-dimensional embedding. Each model followed the original ABB2 protocol (training until no improvement after 100 epochs). This process allowed us to carry out a larger number of experimental runs and only build full models when we had identified the data cutoffs that significantly affected prediction accuracy (assessed by RMSD between predictions and experimental data points).

## Results

### Dominant CDR lengths in paired sequence space are matched by comparable distributions in structural data

We predicted the structures of ∼ 1.5M paired antibody sequences from OAS (Kovaltsuk et al., 2018; Olsen et al., 2022) using ABodyBuilder2 (ABB2) (Abanades et al., 2023), a state of the art deep learning antibody structure predictor. We examined this structural space for evidence of novel canonical forms.

We analysed the length distributions of the CDRL1, CDRL2, CDRL3, CDRH1 and CDRH2 loops in this dataset by both absolute frequency (Fig. 1A) and by non-redundant CDR sequence frequency (Fig. 1B). This revealed that many loops belonging to a specific length were dominated by a smaller number of unique sequences. For example, CDRL1 IMGT length 6 had a frequency of 753,690 in 1.49M, of which only 21,700 were unique. Therefore, we decided to focus our structural analysis on the CDR loops and length combinations (e.g. CDRL3 loops of length 9, after this point referred to as CDRL3-Len-9) which had the highest number of non-redundant sequences within the dataset (bars marked with an asterisk in Fig. 1B, details and numbers given in Table 2).

**Figure 1:**
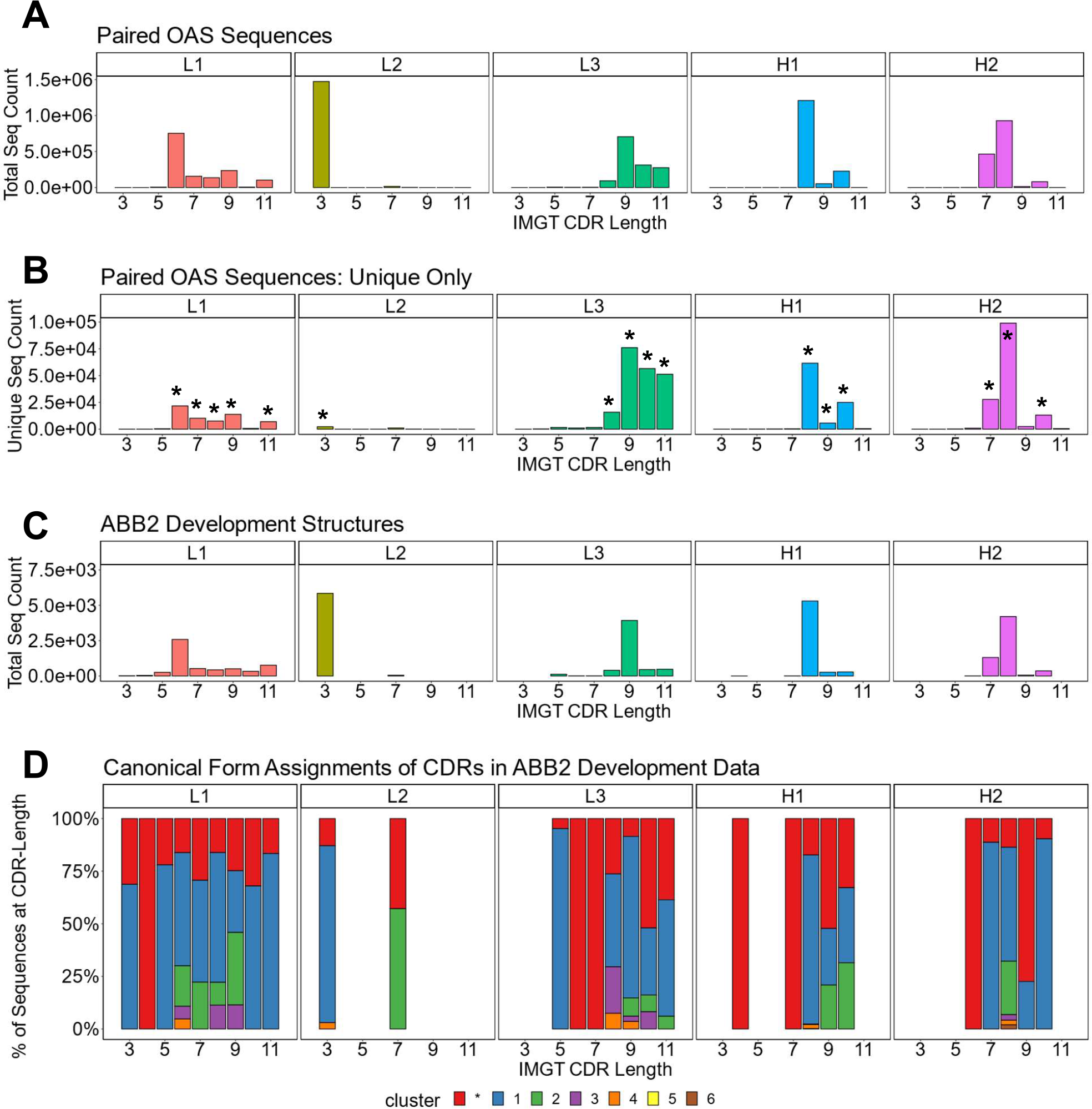
Dominant CDR lengths in paired sequence space are matched by comparable distributions in structural data. Frequency distributions of sequences present in OAS by CDR and loop length, for the total number, including all redundant sequences (A), and unique sequences only (B). Asterisks above certain bars in (B) indicate the lengths with the most unique sequences, the predicted structures of which were analysed. (C) and (D) show data for the antibodies used to develop ABB2. The loop length frequency for each CDR including all structural units (all copies in the asymmetric unit) which have information in PyIgClassify2 (Kelow et al. 2022) are shown (C). Breakdown of canonical form assignments for corresponding CDR loops present in structures of ABB2 training data (D). Each colour within a bar represents a distinct canonical form, red portions indicate loops that could not be assigned to any canonical form with high confidence in PyIgClassify2 analyses (not labelled with a number in the colour legend).

The length distributions of the experimentally derived SAbDab (Dunbar et al., 2014; Schneider et al., 2022) structures used to develop ABB2 are shown in Fig. 1C. Only structures from the train, test and validation datasets which had information on canonical forms detailed in PyIgClassify2 were analysed (Kelow et al., 2022) (38 datasets used in development of ABB2 were not categorised in PyIgClassify2). The CDR loop and length combinations taken forward for further analysis were also enriched in the structural units used to train ABB2 (Fig. 1C), with the minimum number of examples seen by ABB2 during training being 268 for CDRH1-Len-9.

We next analysed the high confidence PyIgClassify2 canonical form assignments of each CDR loop length marked for further investigation by plotting the proportions of each canonical form within all experimental units (Fig. 1D, an experimental unit refers to the fact that one PDB file may have multiple copies in the asymmetric unit). This analysis demonstrated a similar bias, with a single canonical cluster dominating over 50% of assignments for 13 out of the 17 CDR loop and length combinations. Some canonical forms had a very small proportion of examples contained in the ABB2 development data, with the minimum number being 8 examples for CDRH1-Len-8 (Fig. 1D). The biases in both the length and canonical clusters distributions of the experimental data indicate that some data poor areas may benefit from augmentation with predicted structures.

Having selected the CDRs and lengths which dominated our predicted structures, we next built structural clusters for these and explored whether the predicted structures gave rise to new canonical forms or provided insights which were not evident from experimental data alone.

### Predicted structures fall into dense regions of conformational space defined by existing canonical forms

We created a map of the structural space for each of the dominant CDR loop and length combinations, identified above, using the predicted antibody structural data. Each map was analysed to find clusters of CDR loops that shared the same backbone conformation. If a SAbDab structure belonged to a cluster this allowed us to annotate the clusters canonical form according to PyIgClassify2. These annotated maps of canonical form structural space enabled us to navigate the predicted structural space and identify highly occupied regions of space not currently defined by a canonical form.

A full description of how these structural space maps were generated is given in the methods. In brief, each map is a 2D representation of the 3D clustered space of a CDR type at a given length. Data points representing loops from both experimental and predicted structures are coloured by their DBS cluster membership. All data points which are not assigned to a DBS cluster are coloured black. For canonical form annotation the experimental data points are coloured according to their PyIgClassify2 high confidence canonical cluster assignment. Any loops that do not belong to a high confidence PyIgClassify2 cluster (defined by an asterisk in the canonical cluster label) are coloured in red. Figures 2A-B show the structural space map for CDRL1-Len-6. The projections are overlaid with either DBS cluster membership information (Fig. 2A), or canonical cluster classifications from PyIgClassify2 (Fig. 2B). The sequences of the loops which comprised these clusters are visualised using logo plots (Fig. 2C) to identify the motifs and amino acid enrichments which should match to the canonical sequence motifs described in PyIgClassify2 (Table 1). Samples of loops were also inspected in 3D to assess differences in backbone conformations that give rise to the distinct clusters (Fig. 2D).

**Figure 2:**
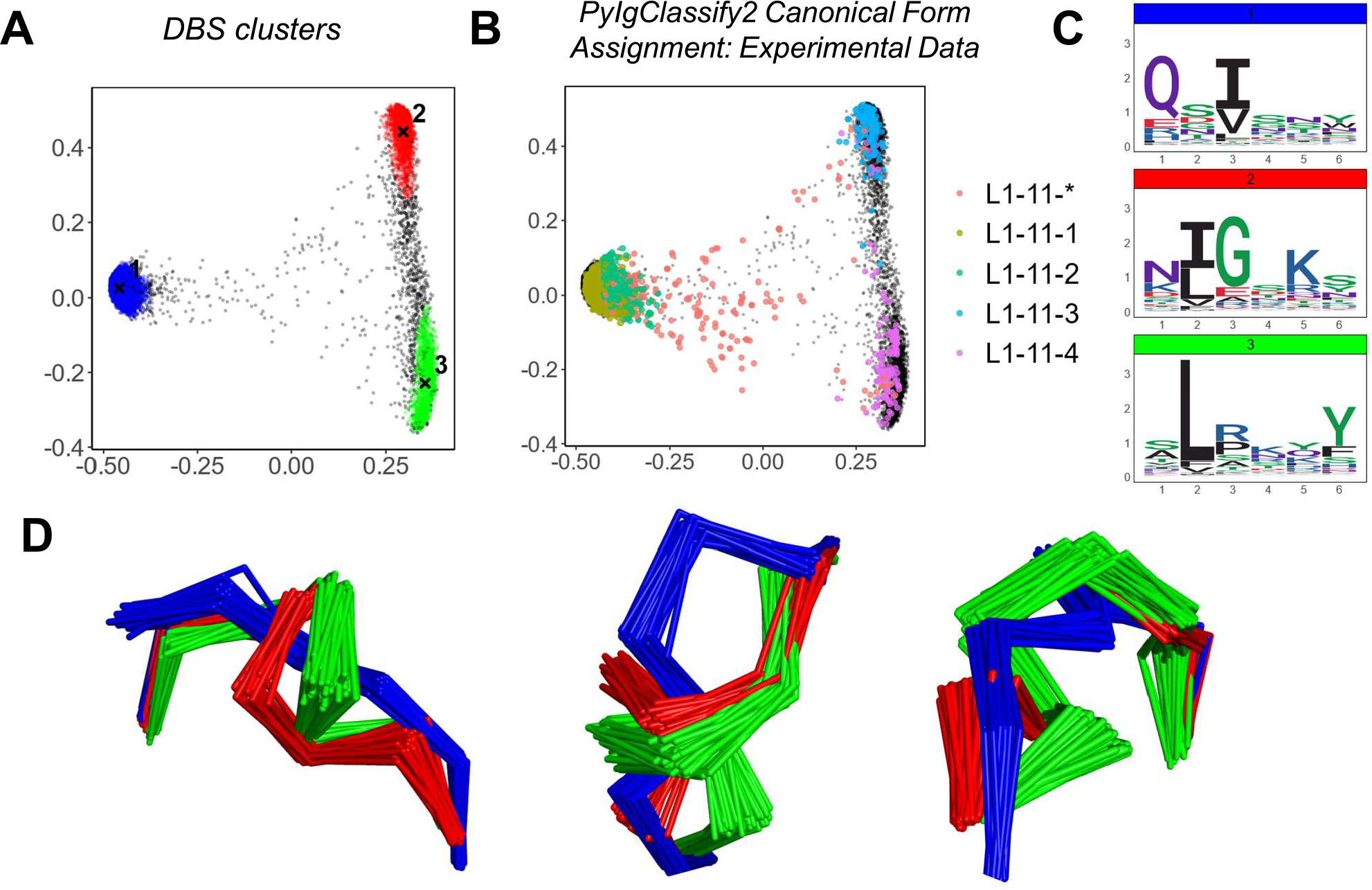
Predicted Structures fall into dense regions of conformational space defined by existing canonical classes. Analysis of CDRL1 length 6 loops. Multi-dimensional scaling plots derived from pairwise RMSD data (A-B). Data points in (A) are coloured according to density-based clustering (DBS) membership, with the cluster centroid marked by an X. Those data points which do not belong to a DBS cluster in (A) are coloured black. In (B) all experimental data points present in the MDS analysis are coloured according to their high confidence PyIgClassify2 canonical form annotation. Any loops that do not belong to a high confidence PyIgClassify2 cluster (defined by an asterisk in the label, e.g. L1-11-*) are coloured in red. The predicted data points are coloured in black and underly the experimental annotations. Logo plots are shown for all sequences in each DBS cluster (C). For a sample of 20 loops in each cluster, the framework aligned backbone conformations are shown in three different orientations (D). The logo plots (C) and backbone (D) colours of blue, green and red correspond to the numbered DBS cluster colours in (A).

For many of the CDR and length combinations analysed in this way, the clusters arising from predicted structures aligned well with the dominant clusters of experimentally defined canonical forms and did not highlight any new areas of density without canonical cluster assignment (Fig. 2, Supp Fig. 1, Table 2). Inspection of loop alignments revealed that our RMSD and DBS analysis method could distinguish between backbone kinks, peptide flips and minor variations that equated to less than 1 Å RMSD between data points (Supp Fig. 2A-C). These minor differences in conformation were also detected by the dihedral angle metric used to compile PyIgClassify2 clusters resulting in similar global divisions of structural space. Before exploring areas that were not accounted for by existing definitions of canonical forms, we next inspected any inconsistencies between our RMSD/DBS analysis and PyIgClassify2.

### Differences between PyIgClassify2 Definitions and Density Based Structural Clusters

While most DBS clusters detected in our analysis could be mapped to experimental data points that adhered to a high confidence canonical form, several of the more subtle PyIgClassify2 definitions were assimilated into a single DBS cluster. We investigated these assimilated data points to assess whether our method was missing important conformational differences.

The loop which best exemplified this was CDRL3Len-8 (Supp Fig. 1A). Our analysis pipeline identified two DBS clusters, the centroids of which were 1.45 Å apart and had distinct sequence motifs (Supp Fig. 1A). Inspection of the PyIgClassify2 canonical clusters demonstrated that cluster 1 (coloured blue in Supp Fig. 1A) defined by the logo motif of QQYysxxT was subdivided into two canonical clusters, of which one had a proline at position 7 (see Table 1, canonical forms L3-8-1 and L3-8-3). We reran our DBS clustering method using an alternate min points term (square root of N data points, opposed to square root of N/2) and found this was able to subdivide the major cluster into two distinct clusters (Supp Fig. 2E) distinguished by the proline at position 7 (Supp Fig. 2F). These two clusters are only 0.4 Å apart and exhibited a large degree of overlap in conformational space (Supp Fig. 2G).

We found additional examples of this within the clusters of loops for CDRL1-Len-6 (Fig. 2B) and CDRH2-Len-8 (Supp Fig. 1C) where the effect was apparent from the smaller number of DBS clusters annotated by a larger number of canonical cluster labels. However, these were often canonical forms assigned to a comparatively low proportion of experimental data points, with more dominant canonical clusters showing greater overlap with our analyses (see later figures). We reasoned that such subtle shifts in conformation would not serve as strong evidence of extrapolation, despite being valid definitions of canonical forms from the dihedral angle perspective. Therefore, their detection was not crucial to our exploratory search for new knowledge arising from structure prediction tools.

### Predicted structures enrich the experimental landscape revealing subdivisions of existing classes with defined sequence motifs

Having confirmed that our clustering pipeline was able to pick out major differences in loop conformations across large datasets, we next investigated the structural clusters and sequence logos which did not sit within with PyIgClassify2 defined canonical forms. These ambiguous clusters could be divided into two categories, those which contained experimental data points defined by a canonical form but could be further subdivided into new clusters with distinct sequence motifs and loop conformations, and those which contained experimental data points that were not assigned to a canonical form.

Firstly, subdivisions of existing canonical clusters which arise from the enrichment of the existing structural space with predicted structures. For example, in CDRH1-Len-8 (Fig. 3A-B) and CDRH1-Len-10 (Fig. 3C-D) as well as CDRL1-Len-11 and CDRH2-Len-7 (Supp Fig. 3A-B), where DBS clusters with distinct amino acid motifs and loop conformations were resolved from the increased structural dataset. The four sub clusters observed in CDRH1-Len-8 (Fig. 3A) and derived from the PyIgClassify2 canonical form H1-13-5 (Fig. 3B, for motifs and length comparisons see Table 1) showed different amino acid patterns at position 2, 4 and 5 of the loop, and had conformations which differed by RMSD of between 0.79-1.43 Å for cluster centroids. While two sub clusters identified from the canonical form H1-15-2 (Fig. 3D, Table 1) in CDRH1-Len-10 loops differed by 1.09 Å and had sequence patterns that differed at six of the ten positions (Fig. 3C).

**Figure 3:**
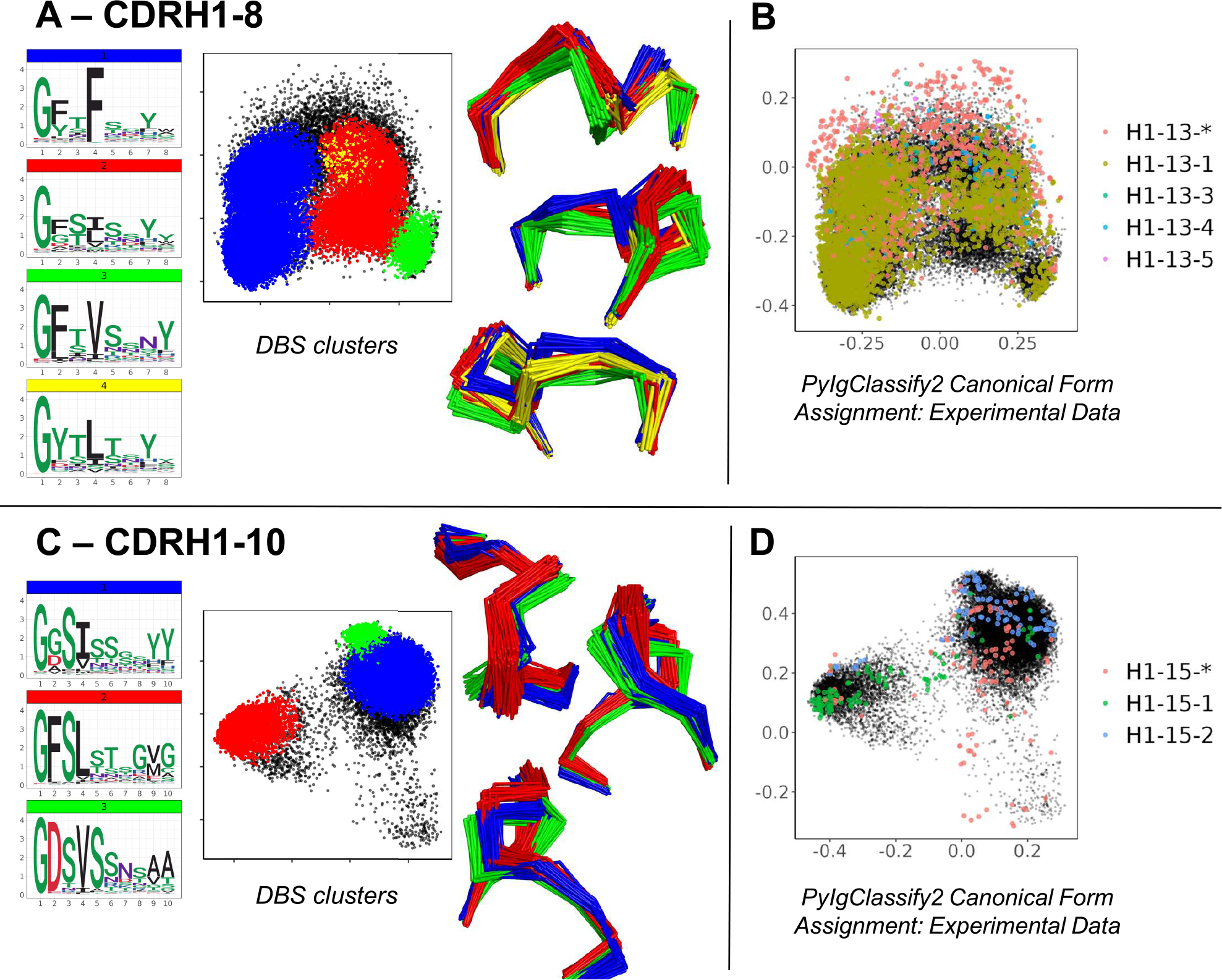
Subdivisions of existing canonical classes into clusters with distinct sequence motifs and backbone conformations. For CDR loops of CDRH1 length 8 (A-B) and CDRH1 length 10 (C-D), distinct DBS clusters contained multiple experimental data points sharing the same canonical form. Panels (A, C) show colour coded DBS cluster logo plots, MDS plots coloured by DBS cluster membership and framework aligned backbone conformations of a sample of 20 loops from each cluster in three different orientations. Data points in the MDS plots which do not belong to a DBS cluster are coloured black. Panels (B, D) show all experimental data points present in the MDS analysis coloured according to their high confidence PyIgClassify2 canonical form annotation. Any loops that do not belong to a high confidence PyIgClassify2 cluster (defined by an asterisk in the label, e.g., H1-13-* and H-15-*) are coloured in red. The predicted data points are coloured in black and underly the experimental annotations.

For all four CDR loops where predictions gave rise to sub clusters, the RMSD between new cluster centroids were often close to the mean value of each analysis (range of mean values from all pairwise comparisons: 0.96 – 1.03 Å, range of distances between cluster centroids: 0.27 – 1.64 Å) and originated from examples present in the training data. Hence the novel canonical classes arising here did not come from generalisation or extrapolation, but simply by statistical power - the increased sensitivity of density-based clustering on the far larger structural set (number of experimental data units versus predicted structures for CDRL1- Len-11: 768 vs 6,992, CDRH1-Len-8: 5,196 vs 61,617, CDRH1-Len-10: 314 vs 2,500 and CDRH2-Len-7: 1298 vs 27,769).

### Enrichment of unassigned areas of structural space defines new canonical forms within heterogeneous sequences

The second set of novel clusters identified related to dense areas of predicted structural space where a smaller number of experimental structures existed but were defined as “unassigned” to any canonical form in PyIgClassify2. For example, for CDRL1-Len-7 a cluster made up of 909 predicted data points (and 32 experimental data points) was identified which was distinct from the centroid of two existing canonical clusters by RMSD values of 3.94 and 4.31 Å respectively (Fig. 4A). This “new” canonical form has a sequence motif with a strong preference for SGH at positions 1-3 of the loop, in contrast to QSV in both existing forms (see logo plot in Fig. 4A, PyIgClassify2 annotations in Fig. 4B and corresponding motifs in Table1). A second example, CDRL1-Len-9 (Fig. 4C), had fewer experimental structures within that area (13 were present in training data) and comprised of 673 predictions. The central motif of INV at positions 3-5 showed no overlap with the enriched residues at the same positions within the three existing canonical forms (Fig. 4D, see Table1), with RMSD values between the corresponding cluster centroids of 2.99-3.51 Å. Both observations were enabled by increased population of structural space with ABB2 predictions from heterogeneous sequences, however they are not evidence of extrapolation given their origins in the training data.

**Figure 4:**
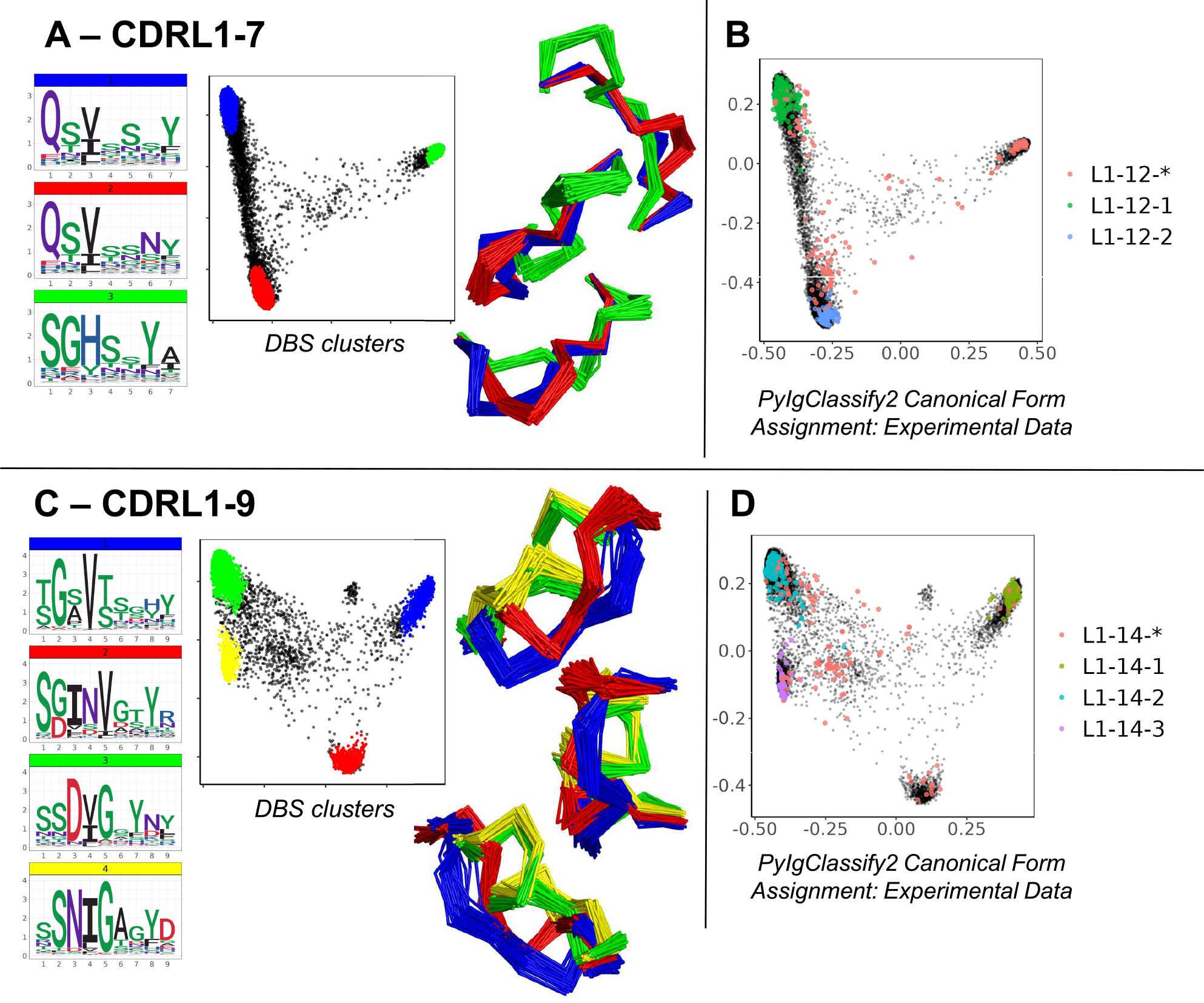
Novel canonical classes revealed by structure prediction. Breakdown of analyses where regions of structural space populated by *unassigned experimental data (denoted by red data points with an asterisk in the PyIgClassify2 annotations) resolved to DBS clusters with distinct sequence logo plots for CDRL1 length 7 (A-B) and CDRL1 length 9 (C-D). Panels (A, C) show colour coded DBS cluster logo plots, MDS plots coloured by DBS cluster membership and framework aligned backbone conformations of a sample of 20 loops from each cluster in three different orientations. Data points in the MDS plots which do not belong to a DBS cluster are coloured black. Panels (B, D) show all experimental data points present in the MDS analysis coloured according to their high confidence PyIgClassify2 canonical form annotation. Any loops that do not belong to a high confidence PyIgClassify2 cluster (defined by an asterisk in the label, e.g., L1-12-* and L-14-*) are coloured in red. The predicted data points are coloured in black and underly the experimental annotations.

### New forms exemplify length independent canonical classes arising from somatic hypermutation

Within the CDRL3 loops we found two further examples of highly populated DBS clusters that did not fit with any PyIgClassify2 definitions. These clusters were identified in the analyses of CDRL3 loops of lengths 10 and 11. Both areas of density contained experimental structures that were classified as unassigned to any high confidence canonical cluster by PyIgClassify2 (CDRL3-10 Fig. 5A-B, and CDRL3-11 Fig. 5D-E).

**Figure 5:**
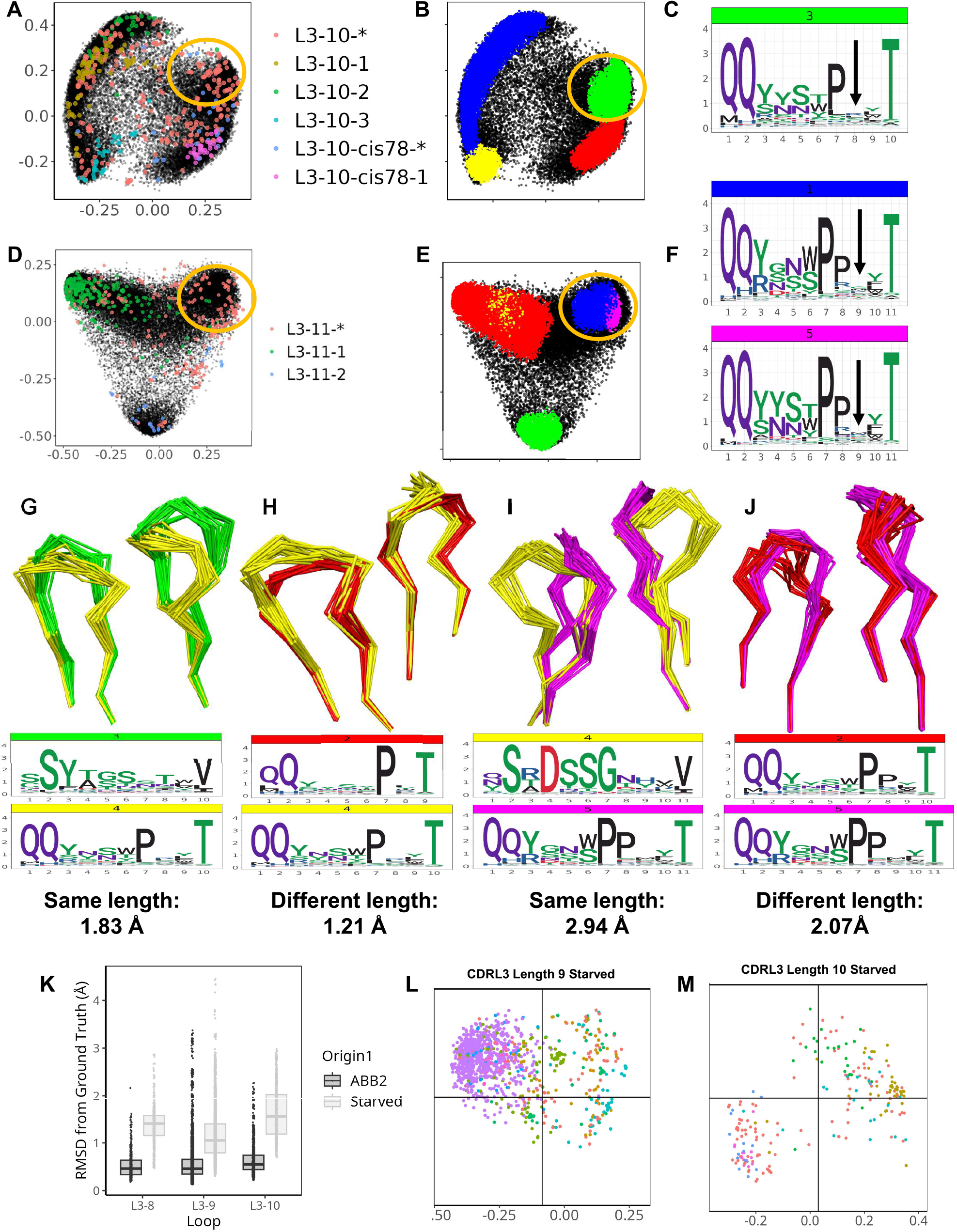
ABB2 can generalise across CDRL3 loops which differ in length by one amino acid. Novel DBS clusters were identified for CDRL3 length 10 (A-C) and CDRL3 length 11 (D-F). Panels (A, D) show experimental data points coloured according to their high confidence PyIgClassify2 canonical form annotation. Any loops that do not belong to a high confidence PyIgClassify2 cluster (defined by an asterisk in the label, e.g. L3-10-* and L3-11-*) are coloured in red. The predicted data points are coloured in black and underly the experimental annotations. Data points in (B, E) are coloured according to density-based clustering (DBS) membership, with the data points which do not belong to a DBS cluster coloured black. The areas circled in yellow on all four MDS plots (A-B, D-E) relate to the DBS clusters of predicted data points, containing experimental data not assigned to any canonical form that were likely to have arisen from length independence. Logo plots were shown with an arrow indicating the high entropy position with no consistent enrichment (C, F) and likely somatic insertion into a shorter canonical cluster (see Supplementary Figure 5). Dynamic time warping analysis permitted quantification of cluster distances between CDR loops of different length as well as clusters of the same length. Clusters of the same length for CDRL3 length 10 are compared by visualising the backbone atoms and sequence logo plots (G), while the novel cluster (coloured yellow) is compared against the proposed origin cluster that the short length of 9 in (H). The DTW distance between cluster centroids is given below each logo plot. CDRL11 clusters of the same length are compared in (I), then the novel cluster and proposed origin cluster in CDRL3 length 10 are compared in (J). Out-of-domain experiments were carried out by retraining ABB2 in the absence of all experimental data points for each of the CDRL3 lengths 8, 9 and 10 (K-M, also see Supplementary Figure 6). Boxplots (median and upper and lower quartiles) and dot plots of RMSD values between predictions and ground truth for each starved model versus the original ABB2 ensemble are compared (K), each dot corresponds to the RMSD value of one comparison. The global conformational space of predictions on withheld data points specific to each model are shown for the CDRL3 length 9 (L) and CDRL3 length 10 (K) starved models. The separation of data points according to canonical form classification was compared to true conformational space and ABB2 ensemble predictions in Supplementary Figure 6 (for each length MDS calculation was performed all data points in the same analysis to allow comparison).

However, these loop shapes are potentially derived from somatic hypermutation (SHM) insertions into CDR loop sequences classified as canonical forms at a shorter length (Supp Fig. 4A-C). These were evident from inspection of logo plots of CDRL3-Len-10 cluster 3 position 8 (Fig. 5C), and CDRL3-Len-11 clusters 1 and 5 at position 9 (Fig. 5F), where the motif was nearly identical to that of a highly populated cluster in the CDR one amino acid length below (PyIgClassify2 canonical forms of L3-9-2 and L-10-cis78-1, see Table 1). The SHM insertions were clearly visible on each logo plot as they resulted in no consensus amino acid enrichment at a fixed position in the loop (positions are marked by an arrow in Fig. 5C and 5F respectively).

Given we could identify the corresponding canonical cluster at the shorter length (we termed this the ‘origin cluster’) (Supp. Fig 4), we decided to quantify and compare the conformation differences between the two sets of loops. To obtain distance scores that could be used for comparison, both for loops of the same length and differing lengths, we substituted pairwise RMSD calculations with dynamic time warping (DTW) calculations which can be performed on coordinate arrays of differing dimensions (see methods for details).

We represented the structural relationships between CDRL3 loops of length 9 and 10 (Supp Fig. 5A), and CDRL3 loops of length 10 and 11 (Supp Fig. 5B) using DTW. The cluster of loops representing the novel conformation in CDRL3-10 (cluster 4: QQYxxxPxxT, coloured yellow in Supp Fig. 5A) was closest in 3D space (DTW distance between cluster centroids of 0.68 Å) to a cluster composed of CDRL3-Len-9 loops (cluster 1: QQYysxxxT, coloured blue in Supp Fig. 5A), and only 1.21 Å away from the proposed origin cluster (CDRL3-Len-9 cluster 2: QQyxxxPxT, coloured red in Supp Fig. 5A). These distances were less than, or comparable to both clusters present at the same length of 10 (distances of 1.83 Å and 1.12 Å apart respectively). The same effect was more pronounced for the novel conformation in CDRL3-Len-11 (cluster 5: QQYxxxPPxxT, coloured pink in Supp Fig. 5B). Here the centroids of clusters found at the same length were 2.48 Å and 2.94 Å away, while the cluster at the shorter length was only 2.07 Å away. Visual inspection of the loop backbones from different clusters shows how similar conformations of mismatched lengths (Fig. 5H, 5J) can be closer in 3D space than matched lengths from different DBS clusters (Fig. 5G, 5I).

These observations fit with the previously described idea of length independence in canonical forms (Nowak et al., 2016), where the closest partner of a structural cluster, or CDR loop, is another cluster, or loop, present at a different length. We hypothesise that the high frequency of predicted structures derived from heterogenous sequences in OAS altered by SHM, helped to reveal these length independent patterns.

### ABB2 can generalise across CDRL3 loops which differ in length by one amino acid

To explicitly test whether the training and test data points with a specific CDR length influence the predictions of CDR loops of a different length we performed several out-of-domain experiments. These involved modifying the ABB2 training and test data to remove all data points containing CDRL3 loops length of 8, 9 or 10 (one length per experiment). In each case a new instance of the ABB2 model was trained on a reduced dataset. The resulting models were used to make predictions from sequences of the withheld length which could be assessed individually for accuracy, and together for occupancy of structural space.

The first model tested was trained in the absence of all 392 datapoints (referring to all copies in the asymmetric unit of each PDB file) that had CDRL3 length 8 loops. Prediction accuracy was poor (Supp Fig. 6A-B), with median RMSD values (prediction versus ground truth) of 1.41 Å compared to 0.46 Å for the fully trained ABB2 model (Fig. 5K). There was no clear separation of canonical clusters in conformational space, with median values for each form above 1 Å from the ground truth structure (Supp Fig. 6B).

**Figure 6:**
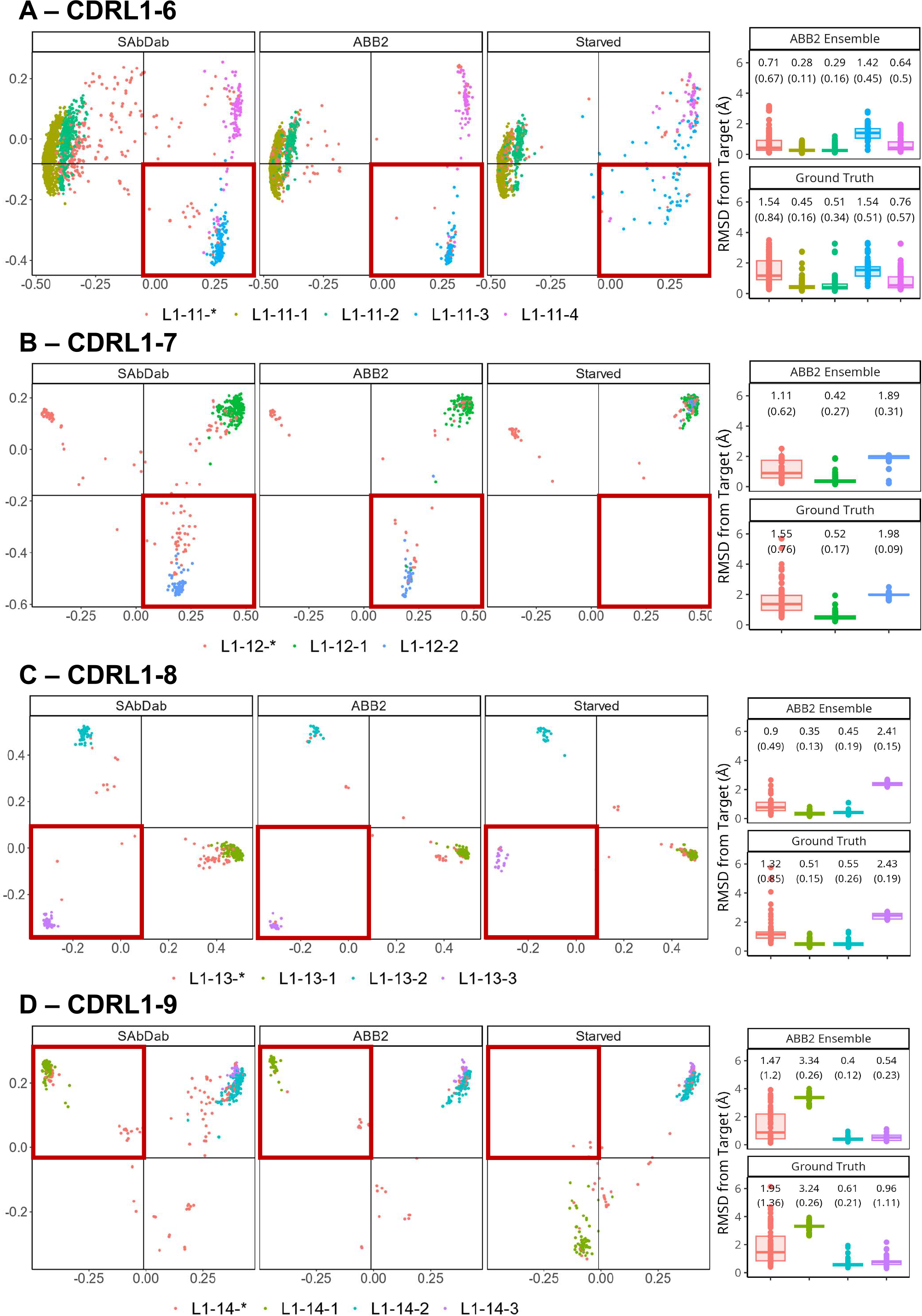
Retraining whilst withholding canonical clusters highlights limited ability to extrapolate. Results of out-of-domain experiments where ABB2 was retrained in the absence of all experimental data points assigned to a specific canonical form in PyIgClassify2 (both high and low confidence) for CDRL1 lengths 6-9 (A-D). MDS plots show experimental data points coloured according to their high confidence PyIgClassify2 canonical form annotation. Any loops that do not belong to a high confidence PyIgClassify2 cluster (defined by an asterisk in the label, e.g., L1-11-*) are coloured in red. CDRL1 length 6 was retrained in the absence of all data points assigned to the PyIgClassify2 cluster ‘L1-11-3’ (coloured blue in panel A). For CDRL1 length 7 ‘L1-12-2’ was dropped (coloured blue in panel B). For CDRL1 length 8 cluster ‘L1-13-3’ was dropped (coloured purple in panel C), and for CDRL1 length 9 cluster ‘L1-14-1’ was dropped (coloured green in panel D). For each panel, the MDS of experimental data points found in SAbDab are shown in the far-left panel. The MDS plot of ABB2 model predictions of the corresponding sequences are shown in the middle panel (labelled ‘ABB2’), and predictions of the starved model are shown in the right panel (labelled ‘Starved’). The area of structural space investigated through exclusion during training is highlighted in a red box. The boxplots (median and upper and lower quartiles) and dot plots on the far right of each panel indicate the RMSD values for each predicted data point from the starved model from its target structure in predictions from the fully trained ABB2 ensemble (top graph) or the ground truth structure (bottom graph) with each sub graph labelled accordingly.

In contrast the model trained in the absence of CDRL3-Len-9 data had better prediction accuracy on the withheld structures (3865 datapoints, median RMSD 1.05 Å) but still worse than the fully trained ABB2 ensemble (median RMSD 0.46 Å). However, the MDS representation of conformational space showed early separation of data points defined by similar canonical clusters (Fig. 5L, Supp Fig. 6C-D), indicating some rationalisation of the sequence to structure relationship via length offset data. Accuracy for the CDRL3-Len-10 model was the worst (median RMSD 1.66 Å Fig. 5K), however a higher standard deviation reflected the correct separation of global conformational space for data points of some canonical forms where loops were close to 1 Å RMSD from the ground truth structure (Fig. 5M, Supp Fig. 6E-F).

There are only 9 data points of CDRL3 length 7, and this may explain why all predictions of CDRL3 length 8 for the starved model fell into a single cluster. In contrast, models starved of CDRL3 lengths 9 and 10 were able to separate some predictions into areas of conformational space close their ground truth structure, this may have been due the abundance of data points either side of the missing length in training data. These experiments, in addition to the structural overlap of predictions of different lengths, provided evidence of generalisation by ABB2 with origins in CDR length independence. Furthermore, these out-of-domain experiments serve as a powerful method to further explore the ability of deep learning-based structure prediction methods to extrapolate and find evidence of truly novel predictions.

### Retraining whilst withholding canonical conformations highlights limited ability to extrapolate

Our analyses so far have not found any evidence of structural clusters representing novel conformations within the CDR loops of predicted antibody structures. Therefore, we set out to explicitly test whether ABB2 could predict a loop conformation not seen in the training data and without a parallel example at a different length. We ran out-of-domain experiments to train models in the absence of all examples of a specific canonical cluster and any close conformations (see methods). We focused on CDRL1 lengths 6-9 as these analyses showed the clearest cluster separation and the smallest proportion of data points that did not fall into a DBS cluster, this helped to avoid any ambiguity in the contents of the training data.

Separate models were trained for each withheld canonical class (numbers and training details given in Table 3). Each model was analysed as before, by predicting the structures of sequences in the withheld data and assessing the individual prediction accuracy as well as total occupancy of structural space.

**Table 3:** Details of retraining in the absence of canonical clusters. Breakdown of the number of antibody variable fragment (FV) structural units used to train ABB2 and the subsequent ‘starved’ models where all units containing a specific canonical form were withheld. The term structural unit is used to account for multiple copies being present in the asymmetric unit of the same PDB file. For each model, the total number of units seen in training is given, followed by the number of removed units. Then the total number of units related to the CDR being withheld is given (for example all units with CDRL1 IMGT length 6), followed by the number of those units remaining after withholding those related to a specific canonical form.

|  | Original Model | CDRL1-6 Starved | CDRL1-7 Starved | CDRL1-8 Starved | CDRL1-9 Starved |
| --- | --- | --- | --- | --- | --- |
| FV structural units seen in training (including all copies in the asymmetric unit) | 5771 | 5459 | 5556 | 5671 | 5554 |
| Dropped units (corresponding to a specific canonical form) | 0 | 312 | 215 | 100 | 217 |
| Total units with CDR and length tested (before withholding a specific form) | - | 2636 | 532 | 428 | 565 |
| Remaining units with CDR tested (after removal) | - | 2324 | 317 | 328 | 348 |

For the ‘starved’ model trained in the absence of CDRL1-Len-6 canonical cluster L1-11-3 (blue dots in Fig. 6A, for sequence motif see Table 1), all predictions failed to match the ground truth conformation (Fig. 6A) with mean (SD) RMSD difference of 1.54 (0.51) Å. The high standard deviation reflects how some predictions were closer (less than 0.5 Å) to the ground truth structure, however the majority adopted a similar conformation to the closest canonical form L1-11-4 (pink dots in Fig. 6A, cluster centroid distance of 1.81 Å) rather than the more distant forms of L1-11-1 or L1-11-2 (green and mustard dots Fig. 6A, combined into one cluster by our method, centroid distance: 2.75 Å). A more pronounced loss of the ability to extrapolate was observed for models starved of a canonical form in the remaining experiments using CDRL1-Len-7 (Fig. 6B) and CDRL1-Len-9 (Fig. 6D). Here all withheld data points fell into a more distant region of conformation space associated with mean prediction accuracies that were much lower, at mean (SD) RMSD values of 1.98 (0.09) Å RMSD for length 7, and 3.34 (0.19) Å for length 9.

For CDRL1-Len-7 the low prediction accuracy may have been due to very similar sequence motifs (DBS cluster: QSVSSSY, corresponding PyIgClassify2 cluster L1-12-1: RASqSVSSSYLa, versus DBS: QSVSSNY, PyIgClassify2 cluster L1-12-2: RASQSVSSNYLA, see Table 1), as well as a small number of examples within the training data (63 examples of the withheld canonical form). In the case of CDRL1-Len-8, the model made predictions of L1-13-3 (purple dots Fig. 6C) that were in a similar region of structural space, however they were still 2.43 (0.19) Å away from ground truth values (Fig. 6C).

These out-of-domain tests clearly demonstrated that the models trained in the absence of a canonical class were unable to recapitulate the correct conformations which could be predicted by the fully trained ABB2 ensemble. The extent of the distance between predictions and ground truth differed for each starved model and related to both sequence similarity as well as structural deviation. This led us to investigate whether prediction accuracy could be recovered by adding very small amounts of training examples back into the model.

### Inclusion of a small number of examples in training is sufficient to recover predictive capacity

We set out to assess whether limited amounts of training data could recover missing predictions and thus quantify the level of representation needed to produce more accurate models. We chose CDRL1-Len-6 and CDRL1-Len-7 as these had the largest standard deviations in prediction accuracy (see box plots Fig. 6). For each we progressively added increasing numbers of data points back into the initial out-of-domain tests resulting in separate models for each addition of data (CDRL1-Len-6 Fig. 7A-B and CDRL1-Len-7 Fig. 7C-D).

**Figure 7:**
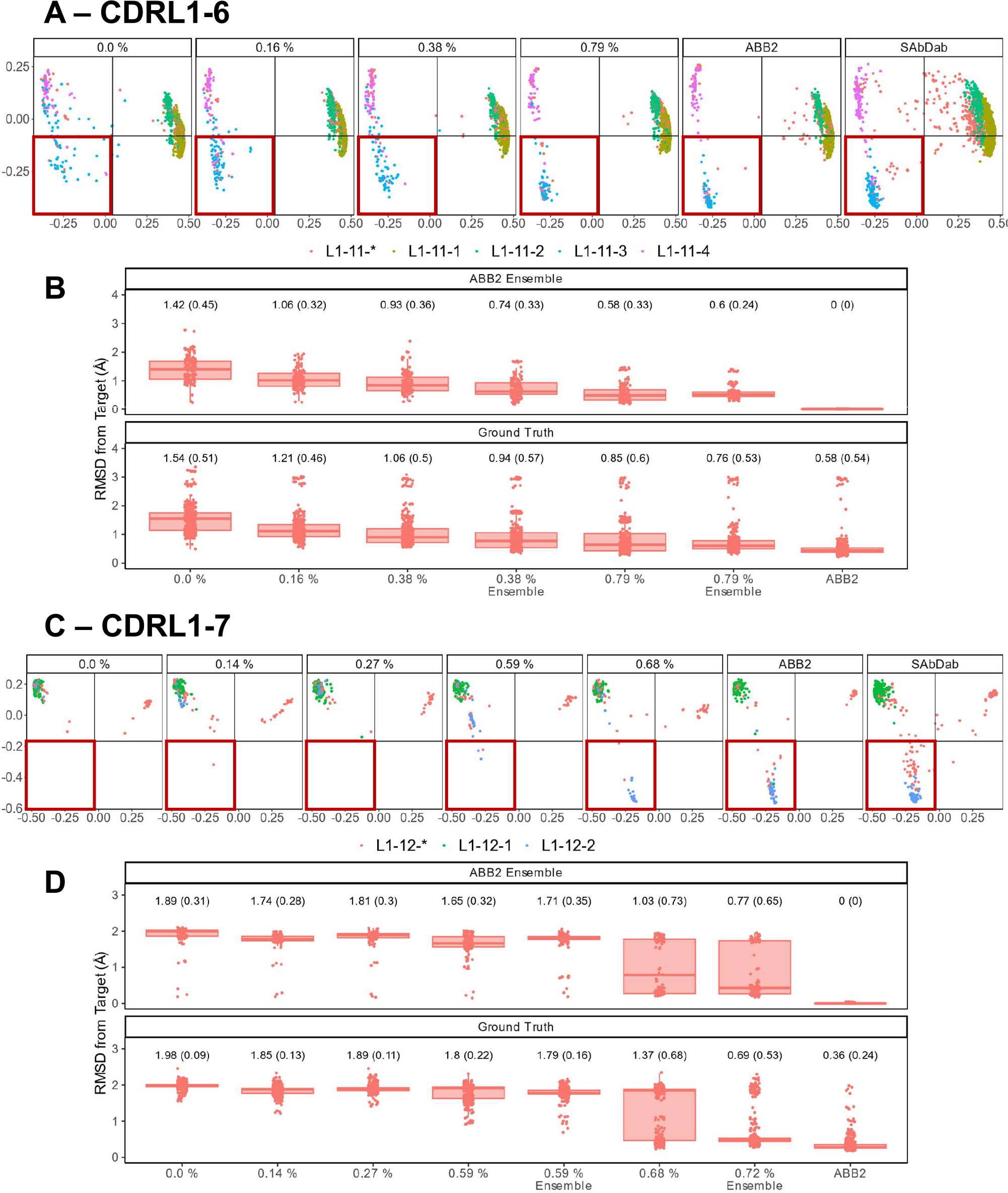
Data points that make up less than 1% of training data are sufficient to recover predictive capacity. Out-of-domain experiments were performed as in Figure 6, however for each model a specified number of experimental data points relating to the withheld canonical form were added into the training data (left to right for each panel, graphs are labelled with the percentages of data points added). Separate models were trained for each of the incrementally increasing data additions until the prediction accuracy came close to the fully trained ABB2 ensemble predictions and the ground truth experimental data. MDS plots for CDRL1 length 6 models (A) and CDRL1 length 7 (C). The amount of data being included is labelled at the top of each panel as a percentage of the total number of unique structures out of all data points seen in training (see Table 4). The corresponding ABB2 predictions and ground truth data are shown in the far-right MDS plots in each panel. RMSD distance values are plotted in (B, D) for the predicted data points relating to the dropped canonical form from each model relative to the ABB2 ensemble predictions (top graph) or ground truth data (bottom graph). For each model, the mean RMSD is given above the box plots, with standard deviation given in brackets. For panels B and D, the top, far-right graph RMSD values are zero as the ABB2 model predictions are being compared to itself, with the comparison of ABB2 to ground truth data shown below.

As there were different total numbers of examples of CDRL1-Len-6 and CDRL1-Len-7 loops in training data (Table 3), the absolute number of PDB structures added back for each experiment did not represent the same proportion of datapoints. Therefore, we calculated the included loops as a percentage of all CDRL1 loops seen for each model (Table 4). Using these values (Table 4) and inspection of the corresponding model performance (Fig. 7), we could see that very small percentages of training data (less than 1%) were enough to allow the models to accurately recapitulate the cluster corresponding to the missing canonical form.

**Table 4:** Inclusion data points as a proportion of total CDRL1 data units in training. For inclusion experiments a specific number of PDB structures were gradually reintroduced to training data for a series of models. As PDB files may contain more than one structure in the asymmetric unit, the number of exact structural units is given for each experiment, as well as the corresponding percentage of all CDRL1 data points present in each training run.

| Number of included PDB structures of withheld canonical form | CDRL1-6 Number of structural units included (percentage of all data points of CDRL1) | CDRL1-7 Number of structural units included (percentage of all data points of CDRL1) |
| --- | --- | --- |
| All | 2636 (45.7%) | 532 (9.2%) |
| 0 | - | - |
| 5 | 9 (0.16%) | 8 (0.14%) |
| 10 | 21 (0.38%) | 15 (0.27%) |
| 19 | - | 33 (0.59%) |
| 20 | 43 (0.79%) | - |
| 21 | - | 38 (0.68%) |
| 22 | - | 40 (0.72%) |

To check that these small proportions of data were enough to facilitate accurate predictions within the original ensemble architecture of ABB2, instead of a single model, we trained ensembles on two inclusion proportions (one above and one below the proportion where we first saw improvement ∼ 0.6%, see Table 4) and analysed the resulting final output (the average structure from all four predictions). This demonstrated that the ensemble models still failed to recapitulate at the lower percentage, while the higher percentages were sufficient for the ensemble to progressively populate the missing structural space despite still being below 1% of total examples of that CDR loop (RMSD plots marked as ‘ensemble’ in Fig. 7B and 7D).

These analyses underline the importance of sufficient data representation and suggest that even small numbers of datapoints can influence the predictive capacity drawn from large datasets. Ultimately the inability of models to truly extrapolate from physical principles means researchers must pay close attention to the contents of their datasets and continue to collect experimental data that explores lesser studied areas of structural space.

## Discussion

In this study we analysed the predicted structures for paired sequences present in OAS generated by ABodyBuilder2 (ABB2) (Abanades et al., 2023). Our data driven approach allowed identification of structural clusters within CDR loops of the same length and subsequent linkage to existing definitions of canonical forms. Our analyses aimed to explore the ability of structure predictors to enrich the experimentally defined landscape of canonical forms and identify novel conformations that would reflect generalisation or extrapolation arising from a deep learning method.

The augmentation of existing data with predicted structures enabled us to define new canonical clusters composed of heterogeneous CDR sequences which were united by the same loop backbone conformation and a sequence motif. These arose from both subdivision of areas of conformational space with uniform PyIgClassify2 annotation, as well as within highly populated areas of conformational space not assigned to any existing canonical form definition. Novel clusters of predicted loop conformations were also produced via the phenomenon of length independence (Nowak et al., 2016). We observed areas of new density which had sequence enrichments identical to those of canonical forms at a shorter CDR length but contained a positionally fixed high entropy residue likely indicative of somatic hypermutation insertion. We analysed the distances between loop conformations at both the shorter and longer lengths by dynamic time warping. This revealed that different length loop clusters were indeed closer in conformational space than any of those of the same length. Out-of-domain experiments confirmed that ABB2 was able to generalise across loops of different lengths and suggested predictions were influenced by high frequency experimental datapoints seen in shorter CDR loops.

However, our analyses could not find new clusters which had no origin in training data and thus represented true extrapolation. Therefore, we performed further out-of-domain experiments by retraining our ABB2 structure predictor whilst withholding data points which belonged to specific canonical forms. These ‘starved’ models were then challenged with correctly predicting the unseen CDR loop conformation. We found that ABB2 retrained in this way was unable to predict conformations not seen during training, but this inability could be resolved by inclusion of a small number of examples representing between 0.5-1.0% of the total data used in development. This suggests that effective prediction accuracy by structure predictors can be achieved for conformations even when they have very poor representation in the dataset.

Our study highlights important limitations regarding the current capabilities of deep learning structure prediction tools specific to the domain of immune receptor CDR loops. Whilst numerous studies have performed out-of-domain experiments and exploratory analysis on protein folds, the conformational space of CDR loops may offer greater challenges, particularly in regions that are inherently flexible or adopt distinct structures in bound and unbound states.

These challenges emphasise that if we wish to predict outside current known structure space, new structure prediction tools that have learnt the underlying rules which govern tertiary structure, instead of just patterns in the training data, will be required. While AlphaFold2 was heralded as a huge advance in structural biology and machine learning, the higher goal of building models that can capture biophysical laws has still not been reached (Buttenschoen et al., 2023; Outeiral et al., 2022). In the absence of architectures that can extrapolate, greater amounts of training data, particularly in regions of structure space with poor coverage, may help improve predictive accuracy. Our demonstration that a small number of examples can address gaps means that a critical mass of data could be achieved to overcome the limitations of current models. However, this does place a large burden on experimental researchers to collect more data.

Finally, our results focused on an area of structural immunology relatively abundant with data and analyses, that of antibodies. As T cell receptors become more important in both immunotherapy research and the clinic, a need to better classify and understand this protein for the purpose of structure prediction may supersede that of antibodies. Therefore, the questions posed in this study may take on more relevance in a field with a relative paucity of structure and paired sequencing data (Leem et al., 2018), as well as several unanswered questions on TCR loop flexibility and comparative conformational freedom (Wong et al., 2019b).

The original purpose of canonical forms was their ability to predict structure from sequence, however these use cases have been superseded by the improved performance of ML methods for structure prediction. For experimental techniques such as X-ray crystallography to be rendered redundant in immunology we must have confidence that structure prediction algorithms faithfully replicate the most important region of immune receptors, the CDR loops.

## Abbreviations

ABB2: ABodyBuilder2
AF2: AlphaFold2
AFM: AlphaFold-Multimer
CDR: complementarity determining region
DBSCAN: density-based spatial clustering of applications with noise,
DBS: DBSCAN-based selection
DTW: dynamic time warping
KNN: K-nearest neighbours
IMGT: the international ImMunoGeneTics information system
MDS: multidimensional scaling
OAS: observed antibody space
OPIG: Oxford protein informatics group
PDB: protein data bank
RMSD: root mean-squared deviation
SAbDab: the structural antibody database
SCFV: single chain variable fragment.

## Contributions

AGW and CD conceived the project and designed the study. BAK contributed to experimental design and data generation. AGW performed experiments and data analysis. AGW, BAK and CD wrote the manuscript. CD supervised the project. All authors contributed to the article and approved the submitted version.

## Acknowledgements

The authors wish to thank Joao Diniz Brandao Gervasio and Carlos Outeiral Rubiera from the Oxford Protein Immunoinformatics Group for their helpful comments and discussions.

## Data availability statement

The structures predicted from Observed Antibody Space have been made available at: https://doi.org/10.5281/zenodo.10280181

