## SupplementaryFigures for "Investigating the ability of deep learning-based structure prediction to extrapolate and/or enrich the set of antibody CDR canonical forms"

### Supp Figure 1: Overlap between PyIgClassify2 canonical form annotations and density-based clusters.

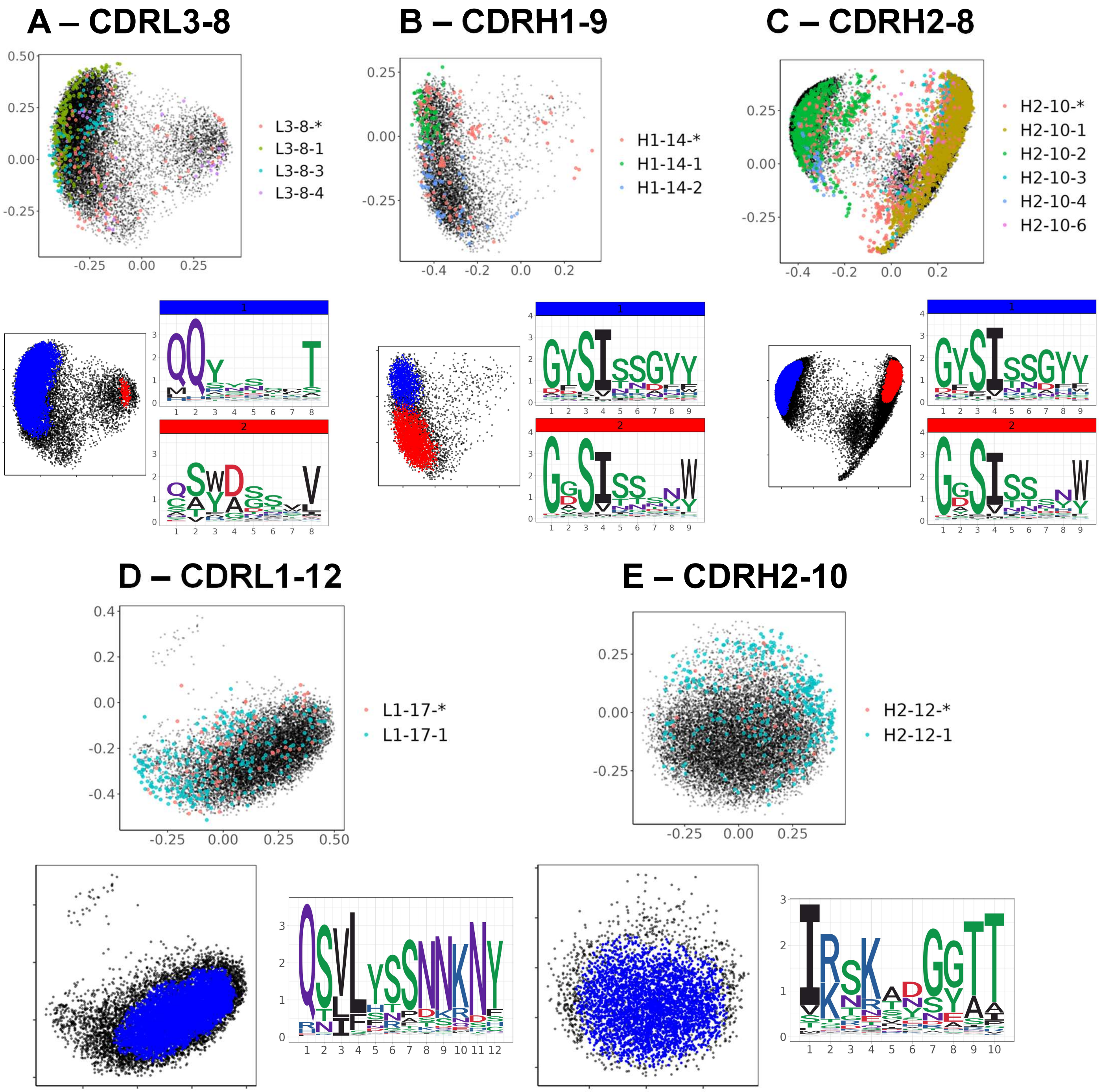


**Supp Figure 1: Overlap between PyIgClassify2 canonical form annotations and density-based clusters.**

For CDRL3 length 8 (**A**), CDRH1 length 9 (**B**) and CDRH2 length 8 (**C**) the clusters of experimental data points and their canonical form annotations by PyIgClassigy2 is shown on MDS plots overlaid on top of the predicted datapoints in black. The corresponding density-based clusters (DBS) are shown on an MDS plot below with the colour coded corresponding sequence logo plots. For each analysis in **A-C** the RMSD and DBS analysis found two dominant clusters which could be related to the PyIgClassify2 defined canonical forms that dominated the data. For CDRL1 length 12 (**D**) and CDRH2 length 10 (**E**), only one DBS cluster was found for each analysis, and this corresponded to a single cluster defined in PyIgClassify2 classifications.

#### Supp Figure 2: Exploration of backbone RMSD differences and minor PyIgClassify2 clusters missed by RMSD and DBS analyses.

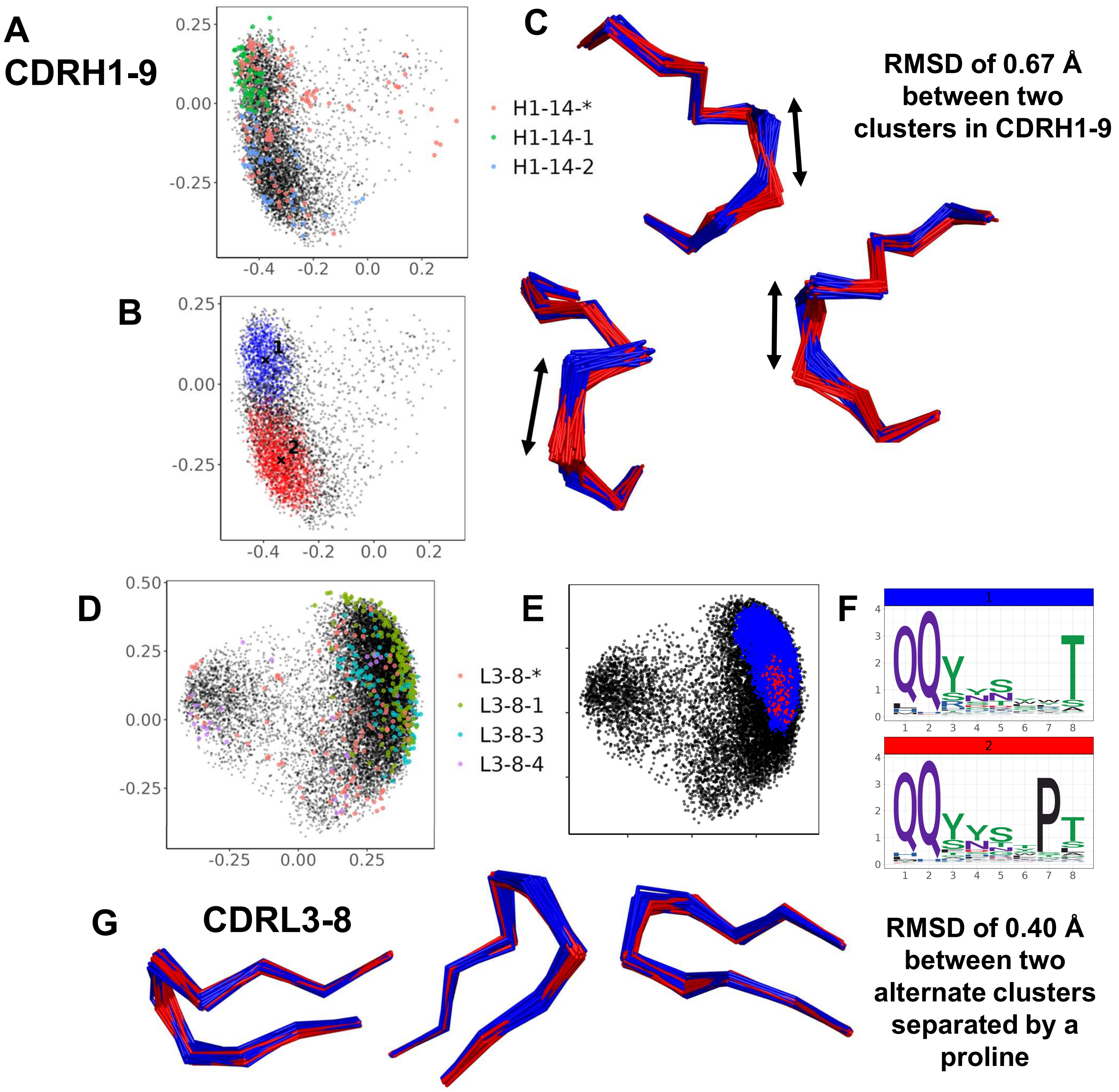


**Supp Figure 2: Exploration of backbone RMSD differences and minor PyIgClassify2 clusters missed by RMSD and DBS analyses.**

For CDRH1 length 9 (**A-C**) the loop backbone RMSD values that lead to distinct DBS clusters were examined. PyIgClassify2 annotations (**A**) and DBS clusters (**B**) were overlaid on MDS plots, as well as backbone visualised from samples of 20 loops belonging to each cluster (**C**) to understand how small shifts in conformation resulted in distinct clusters that related to PyIgClassify2 canonical forms.

For CDRL3 length 8 there was an additional cluster in PyIgClassify2 annotations that was undetected by the initial DBS analysis shown in Supp Fig. 1A, this was ‘L3-8-3’ (**D**) and defined by a proline sequence enrichment at position 7. DBS analysis was rerun with a difference in the min points term (see methods) and this revealed the sub cluster (**E**) with corresponding enriched proline (**F**). Visual inspection of loop backbones demonstrated that despite sequence differences these loops showed a high degree of overlap that made them almost indistinguishable in the absence of colour coding, with an RMSD difference of 0.40 Å between cluster centroids.

#### Supp Figure 3: Subdivisions of existing canonical classes into clusters with distinct sequence motifs and backbone conformations – extra examples.

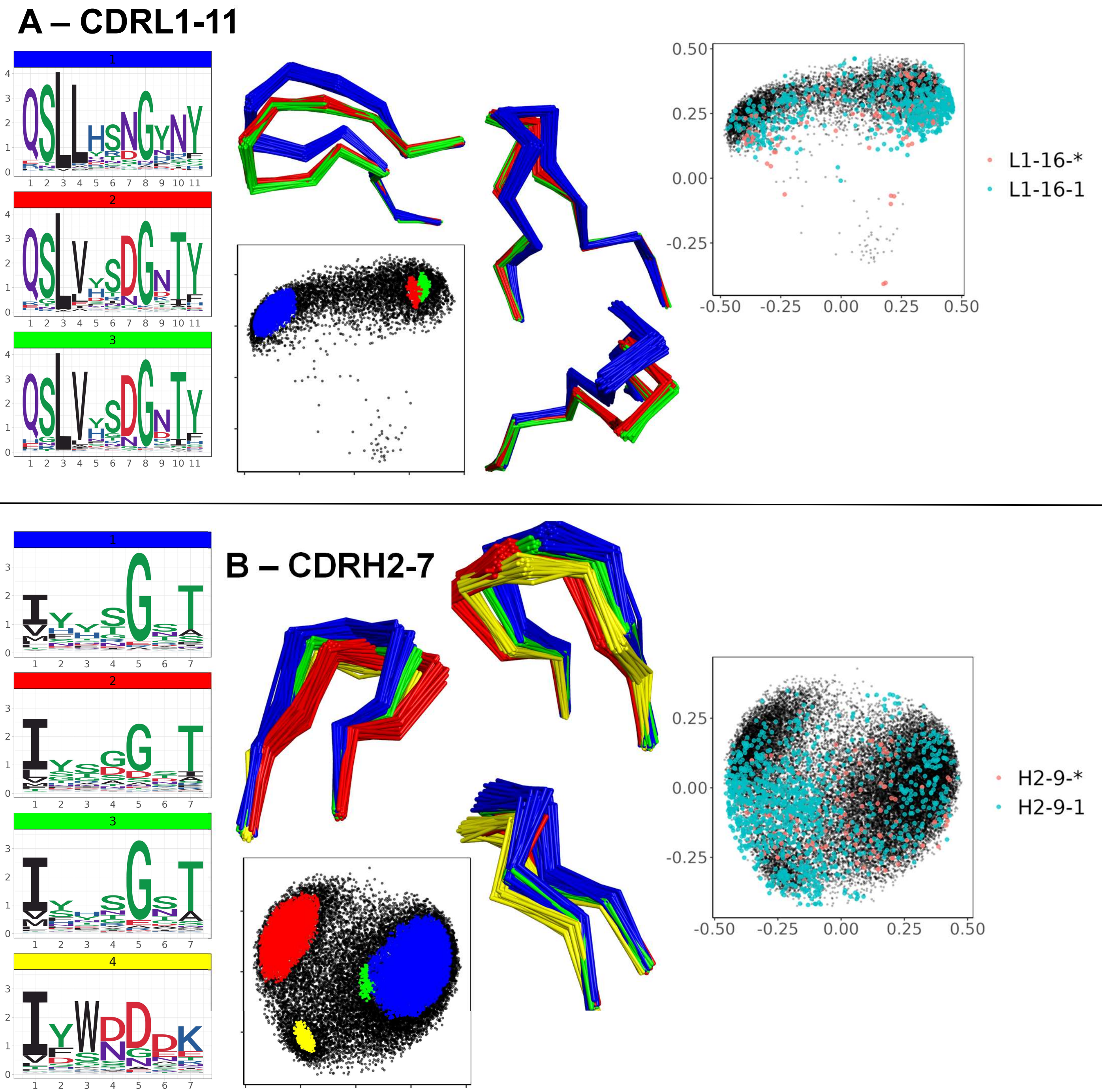


**Supp Figure 3: Subdivisions of existing canonical classes into clusters with distinct sequence motifs and backbone conformations – additional examples.**

For CDR loops of CDRL1 length 11 (**A**) and CDRH2 length 7 (**B**), as shown in Figure 3. Visualisation of distinct DBS clusters which contained multiple experimental data points sharing the same canonical form. Left to right for each panel shows logo plots, an MDS plot annotated by DBS cluster, loop backbone conformations and an MDS with experimental data points coloured by PyIgClassify2 canonical form assignment.

**Supp Figure 4: Novel forms were related to SHM insertions in sequences associated with shorter canonical clusters.**

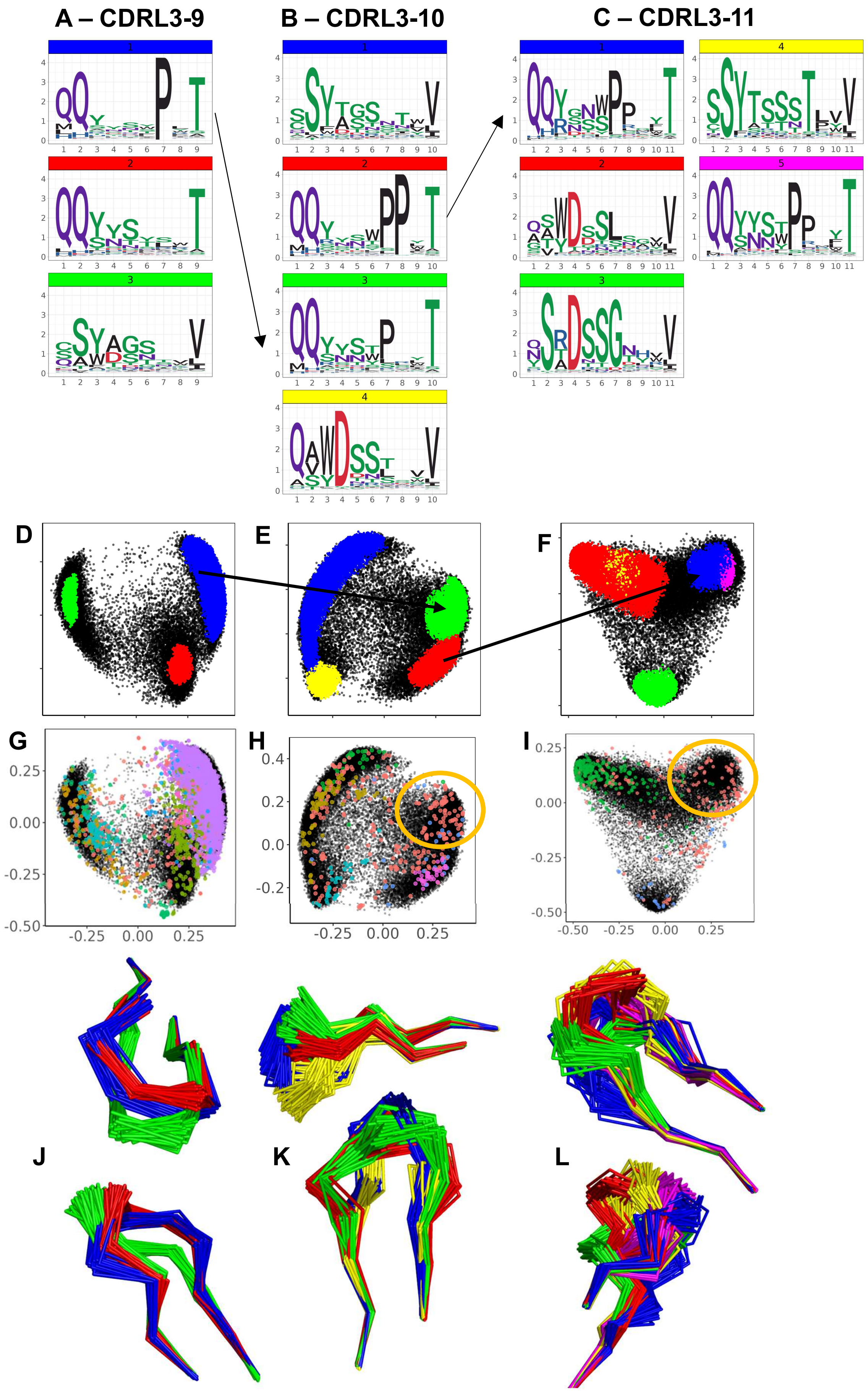

**Supp Figure 4: Novel forms were related to SHM insertions in sequences associated with shorter canonical clusters.**

For CDRL3 loops of length 9, 10 and 11. Sequence logo plots are shown in **A-C** of the corresponding DBS clusters identified, annotated on MDS plots (**D-F**). Between the logo plots and the DBS clusters arrows are shown to indicate the origin cluster at the shorter length into which somatic hyper mutation insertion is hypothesised to have taken place. The corresponding MDS plots with canonical cluster annotations indicate these clusters are annotated with experimental data points classified as unassigned to any canonical form (red datapoints circled with in yellow and marked with an asterisk **G-I**). Corresponding loop backbones are shown in **J-K**.

**Supp Figure 5: Novel forms exemplify length independent patterns**

#### A – CDRL3-9 to CDRL3-10

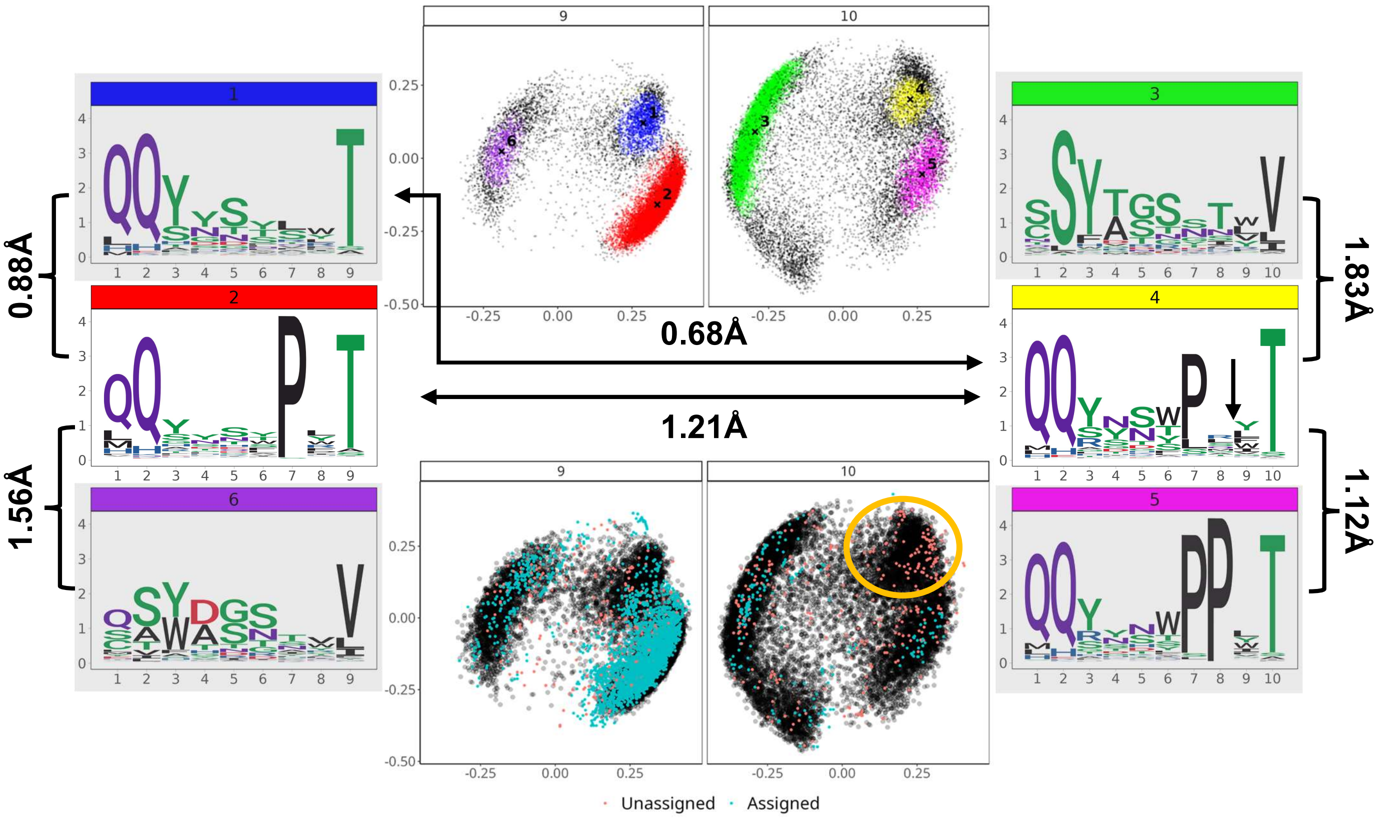


#### B – CDRL3-10 to CDRL3-11

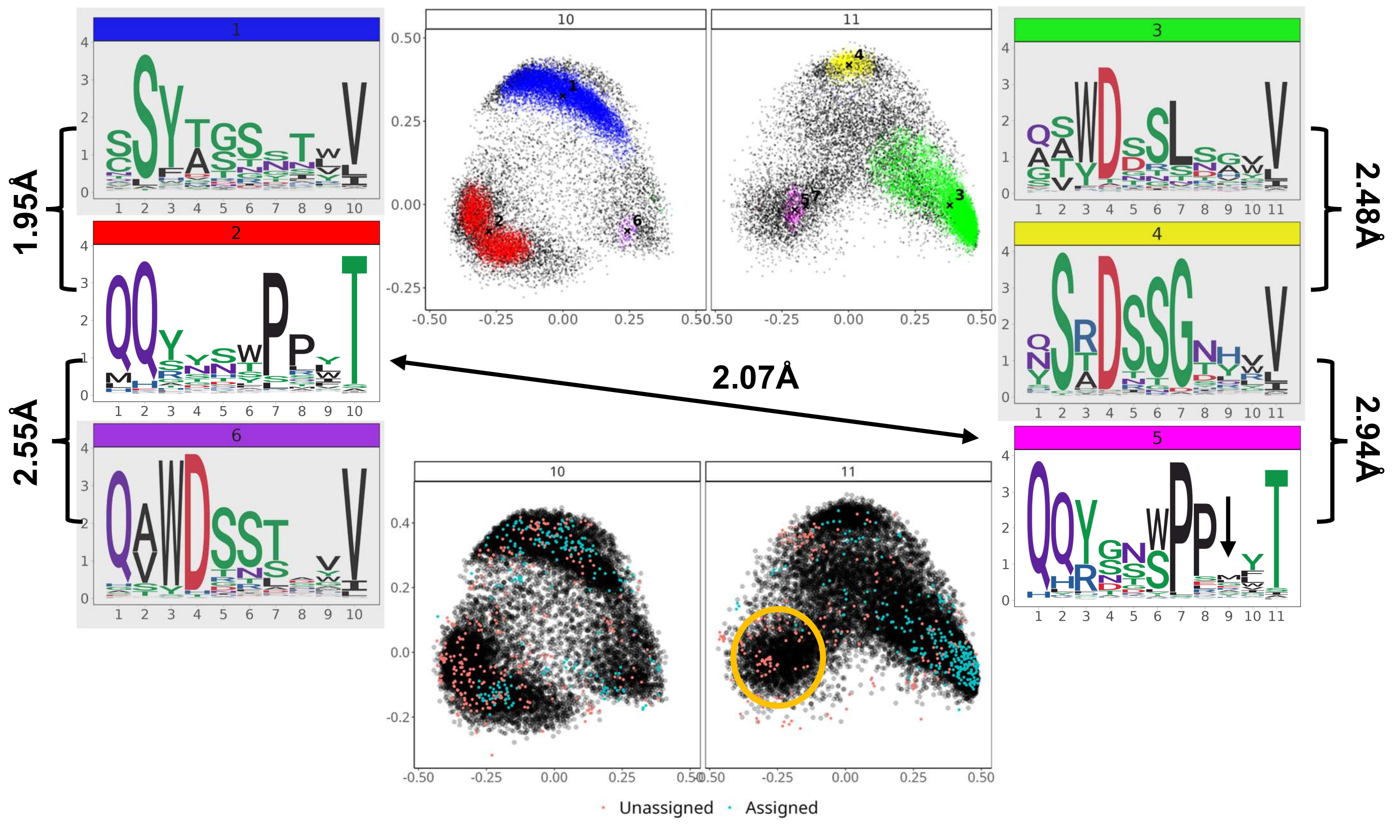

**Supp Figure 5: Novel forms exemplify length independent patterns.**

Dynamic time warping analyses were carried out to compare clusters of CDR loops of different length as well as clusters of the same length. Two analyses were performed CDRL3 length 9 versus length 10 (**A**) and CDRL3 length 10 versus length 11 (**B**). For each analysis, the shorter length logo plots of the DBS clusters are shown on the left, while the logo plots of sequences which comprise the longer clusters are shown on the right.

MDS plots derived from dual length pairwise DTW analyses (and therefore of comparable axes and global position) are faceted according to length, and annotated by DBS cluster in the top plots, or whether they are assigned (cyan) or unassigned (red) to a PyIgclassify2 canonical cluster in the bottom plots. Arrows and connectors indicate the DTW distances between centroids of the DBS clusters and exemplify how clusters of the same length can be more distance than those of distinct lengths.

The novel clusters identified in Figure 5 and Supp Fig. 5 are also circled in yellow in the MDS plots. The logo plots of these novel cluster derived from length independence are shown to the right (not grey scaled), they are coloured by the yellow bar for CDRL3 length 10 (**A**) and pink bar for CDRL3 length 11 (**B**). The corresponding origin clusters are shown on the left for the shorter length and not grey scaled.

### Supp Figure 6: Models trained in the absence of CDRL3 length 8, 9 and 10 data.

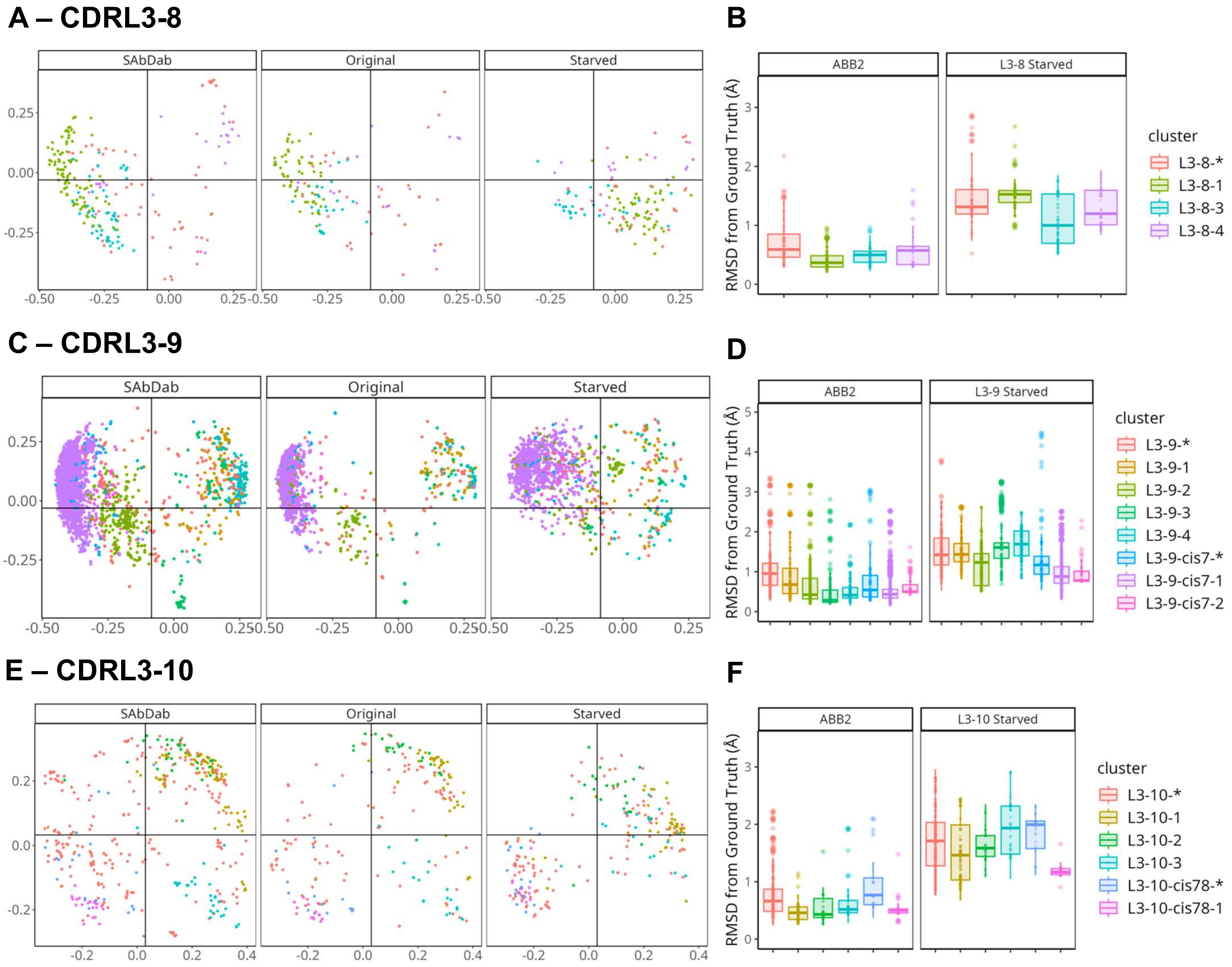


**Supp Figure 6: Models trained in the absence of CDRL3 length 8, 9 and 10 data.**

Out-of-domain experiments were performed by removing all datapoints with CDRL3 length 8 (**A, B**), length 9 (**C, D**) and length 10 (**E, F**). The MDS plots of the ground truth data (labelled 'SAbDab' in the far-left plot), the ABB2 ensemble predictions (middle plot labelled 'Original') are compared to the predictions from the starved models in the far-right plot (**A, C, E**).

Plots in **B**, **D**, and **F** show the RMSD difference of predictions from ground truth experimental data points. The differences between ABB2 ensemble predictions and ground truth are shown in the left plots, while the difference between starved model predictions and ground truth data are shown in the right plots.

**Supp Figure 7: Clustering analysis on batches of structures where number of non-redundant sequences were greater than 42,000.**

#### A – CDRL3-9

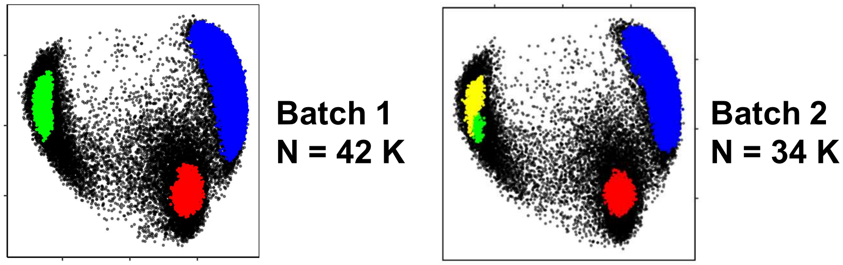


#### B – CDRL3-10

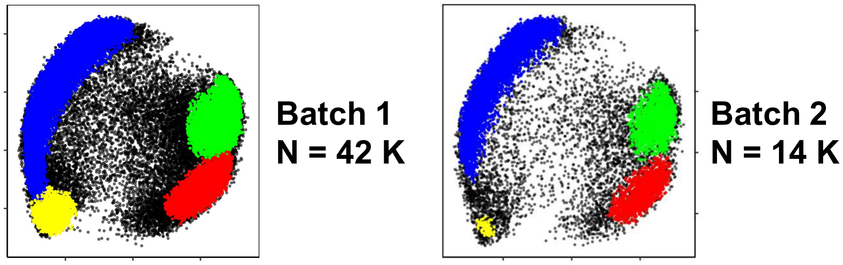


#### C – CDRL3-11

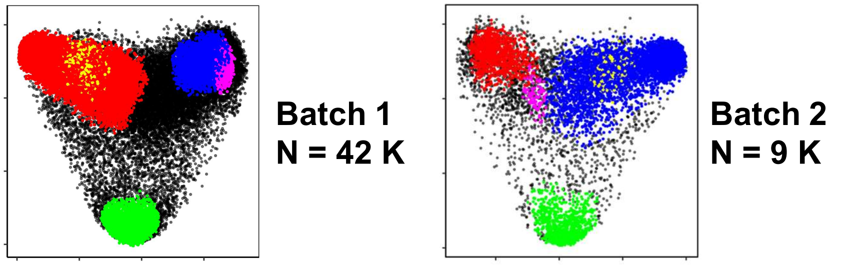


#### D – CDRH1-8

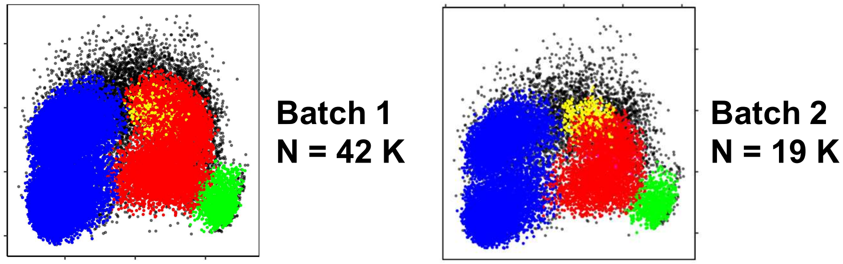


#### E – CDRH2-8

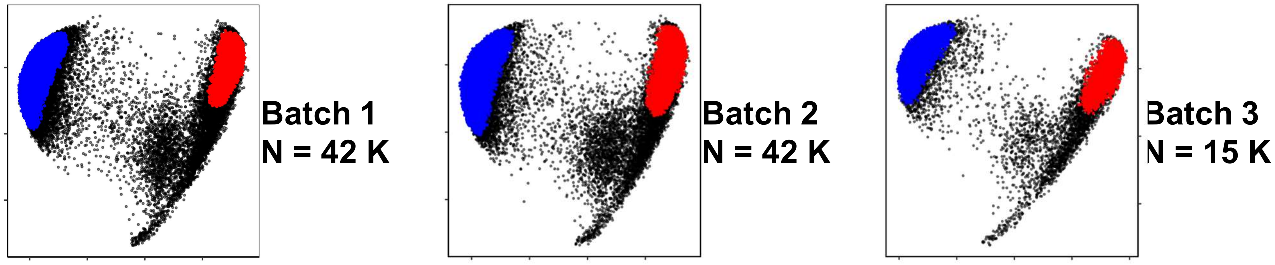

**Supp Figure 7: Clustering analysis on batches of structures where number of non-redundant sequences were greater than 42,000.**

CDR loops of non-redundant sequence identity were analyzed by pairwise RMSD and density-based clustering in batches of 42,000. The CDR and lengths combinations shown in (**A**-**E**) contained more than 42,000 non redundant sequences corresponding to loops within predicted structures. These loops were run through the analysis pipeline in separate batches to ensure every data point was represented and the resulting clusters from each set of samples did not differ greatly between batches. Batches were run with the maximum number of loops first (42,000) and the remaining loops used in subsequent batches. The numbers analyzed in each batch are shown beside each MDS plot.
